# A neuroprosthetic system to restore neuronal communication in modular networks

**DOI:** 10.1101/514836

**Authors:** S. Buccelli, Y. Bornat, I. Colombi, M. Ambroise, L. Martines, V. Pasquale, M. Bisio, J. Tessadori, P. Nowak, F. Grassia, A. Averna, M. Tedesco, P. Bonifazi, F. Difato, P. Massobrio, T. Levi, M. Chiappalone

**Author notes:** Corresponding authors Dr. Michela Chiappalone, PhD Rehab Technologies IIT-INAIL Lab, Istituto Italiano Di Tecnologia, Via Morego 30, 16163 Genova Prof. Timothée Levi, PhD Laboratoire de l’Intégration du Matériau au Système (IMS), University of Bordeaux, Bordeaux INP, CNRS UMR 5218, 351 Cours de la Libération, 33405 Talence Cedex. First Equal (S. Buccelli, Y. Bornat). Second Equal (I. Colombi, M. Ambroise, L. Martines).

## Abstract

Recent advances in neurotechnology allow neurological impairments to be treated or reduced by brain machine interfaces and neuroprostheses. To develop energy-efficient and real-time capable devices, neuromorphic computing systems are envisaged as the core of next-generation ‘neurobiohybrid’ systems for brain repair. We demonstrate here the first exploitation of a neuromorphic prosthesis to restore bidirectional interactions between two neuronal populations, even when one is damaged or completely missing. We used in vitro modular cell cultures to mimic the mutual interaction between neuronal assemblies and created a focal lesion to functionally disconnect the two populations. Then, we employed our neuromorphic prosthesis for two specific applications with future clinical implications: *bidirectional bridging* to artificially reconnect two disconnected neuronal modules and *hybrid bidirectional bridging* to replace the activity of one module with a neuromorphic spiking neural network. Our neuroprosthetic system opens up new avenues for the development of novel bioelectrical therapeutics for human applications.

## Introduction

One of the greatest challenges of modern neuroscience is to find reliable and sustainable treatments for the disabling effects caused by many chronic and incurable brain conditions. With the greatest impact carried by stroke (*1*) and traumatic brain injury (*2*), brain disorders are among the leading causes of disabilities worldwide. Due to recent advances in neural and neuromorphic engineering, direct interfacing of artificial circuits with large neuronal networks is possible to develop novel ‘neurobiohybrid’ systems (such as neuroprostheses (*3*)), which are envisaged as potentially interesting clinical applications for brain lesions (*4*). In this paper, we introduce an innovative neuroprosthetic system based on a neuromorphic real-time interface that can re-establish the communication between two disconnected neuronal assemblies.

Neural interfaces are promising solutions for brain repair (*5*). Modern neural interfaces are mainly designed to restore lost motor functions in only one direction, i.e., from the brain to the body (*6*) or from the body to the brain (*7*). Additionally, recent neuroprosthetic developments have shown the enormous potential of neural interfaces to aid and accelerate functional recovery (*8, 9*). However, a major obstacle in developing novel neuroprosthetic devices for bidirectional communication with and within the brain is the complex nature of interactions among different brain areas, which in turn presents a challenge for the development of appropriate stimulation protocols as well as for testing such devices using in vivo models (*10*).

Despite very recent technological progress (*11, 12*), in vivo models still have two main bottlenecks. The first bottleneck is the technical challenge to faithfully reproduce specific/focal network lesions (mainly due to their complexity) that the neuroprosthesis aims to treat, whereas the second is the difficulty in disentangling the actual effect of the adopted electrical therapy from the complex activity of a brain constantly processing sensory inputs and producing behavior. Therefore, since in vivo models exhibit inherent complexity and low controllability, using in vitro reduced neuronal systems to model precise and reproducible neuronal network lesions and test neuroprosthetic devices for their treatment may be advantageous. This approach is also justified by a growing recognition that in vitro testing of both research and medical devices can be more effective in terms of cost, time consumption and ethical issues and much more reliable than in vivo testing (*13*). Therefore, in this work, we used an in vitro system based on interconnected ‘neuronal assemblies’ (*14*), which can be interfaced bidirectionally in real time with a neuroprosthetic system. Based on our previous experience, we developed bimodular neuronal networks on micro-electrode arrays (MEAs) to record electrophysiological activity and deliver electrical patterns of stimulation (*15*)

Therefore, our first objective was to create a simplified yet plausible in vitro model of a focal brain lesion by using bimodular cultures (*15-17*) with reciprocal connections cut by a custom-made laser setup (*18*) to mimic the pathological effect of TBI (*19*). We created a neurobiohybrid system connecting the biological element (modular culture) following the lesion with a neuroprosthetic prototype. Our neuroprosthetic device can perform low-power computations in hard real time (*20*), collecting the inputs coming from neural recordings and generating suitable electrical stimulation triggers as an output. With this experimental setup, we tested two specific applications exploitable for clinical purposes, namely, *bidirectional bridging* (BB) to artificially reconnect two disconnected neuronal modules and *hybrid bidirectional bridging* in which a neuromorphic spiking neural network (SNN) replaced the activity of one of the two modules in real time while implementing bidirectional connectivity with the remaining neuronal module.

The motivation of our research is to provide a new technological instrument as a novel form of neuroprosthesis aimed at treating disabling brain pathologies. The adoption of bidirectional communication allows the development of a generalized non-specific approach that is applicable to the central nervous system (CNS) or peripheral nervous system (PNS). Indeed, prosthetic systems for the PNS restore the communication between a neuronal assembly and sensor/actuator, and the communication follows a well-defined anatomical and functional path (from neuronal assembly to actuator or from sensor to neuronal assembly). Moreover, prostheses for the CNS should restore the communication between two or more neuronal assemblies whose functional and anatomical path could be distributed and sparse and not necessarily known *a priori*.

Indeed, our idea to develop a generalized approach comes from the future perspective of creating a cerebral neuroprosthesis for direct implantation in the brain that could be used by patients affected by stroke or brain injury. Our proof-of-principle results are the first for a next-generation neurobiohybrid system to restore brain functions (*3, 4*).

## Results

### Neuroprosthetic System

To create bimodular in vitro systems, we developed PDMS masks with two connecting compartments that constrained the growth of neuronal cells in two precise areas over 60-electrode MEAs (Fig. S1). The obtained bimodular neuronal culture constitutes the biological neuronal network (BNN) of our system (Fig. 1A). The signal from the BNN was amplified by a commercial system and acquired by a custom-developed neuromorphic board (Fig. S2) based on Field Programmable Gate Array (FPGA) previously configured by a custom-made MATLAB code (MathWorks, Natick, MA, USA) running on a general purpose personal computer. The neuromorphic board triggered a commercial stimulator to close the loop with the BNN. The general protocol designed for this study involved three steps. First, spontaneous activity in both neuronal modules was recorded (‘pre-lesion condition’). Then, laser ablation of the biological connections between the two modules was performed (see Methods), followed by recording of spontaneous activity in both modules (‘post-lesion condition’) to assess the viability of the networks. Finally, we tested our neuroprosthetic device using two experimental frameworks. In the first case, we applied a reconnection strategy using a bidirectional activity-dependent stimulation (‘bidirectional bridging’, BB), whereas in the second case, we interfaced a hardware-implemented biomimetic SNN with one of the two neuronal modules (‘hybrid bidirectional bridging’, HBB) to simulate a ‘replacement’ strategy that utilizes the bidirectional interaction between the biological system and its artificial counterpart (Fig. 1B). The sequence of algorithms (e.g., spike detection, network burst detection) implemented on the board to realize both experimental approaches (BB or HBB) is schematically depicted in Fig. 1C.

**Fig. 1.**
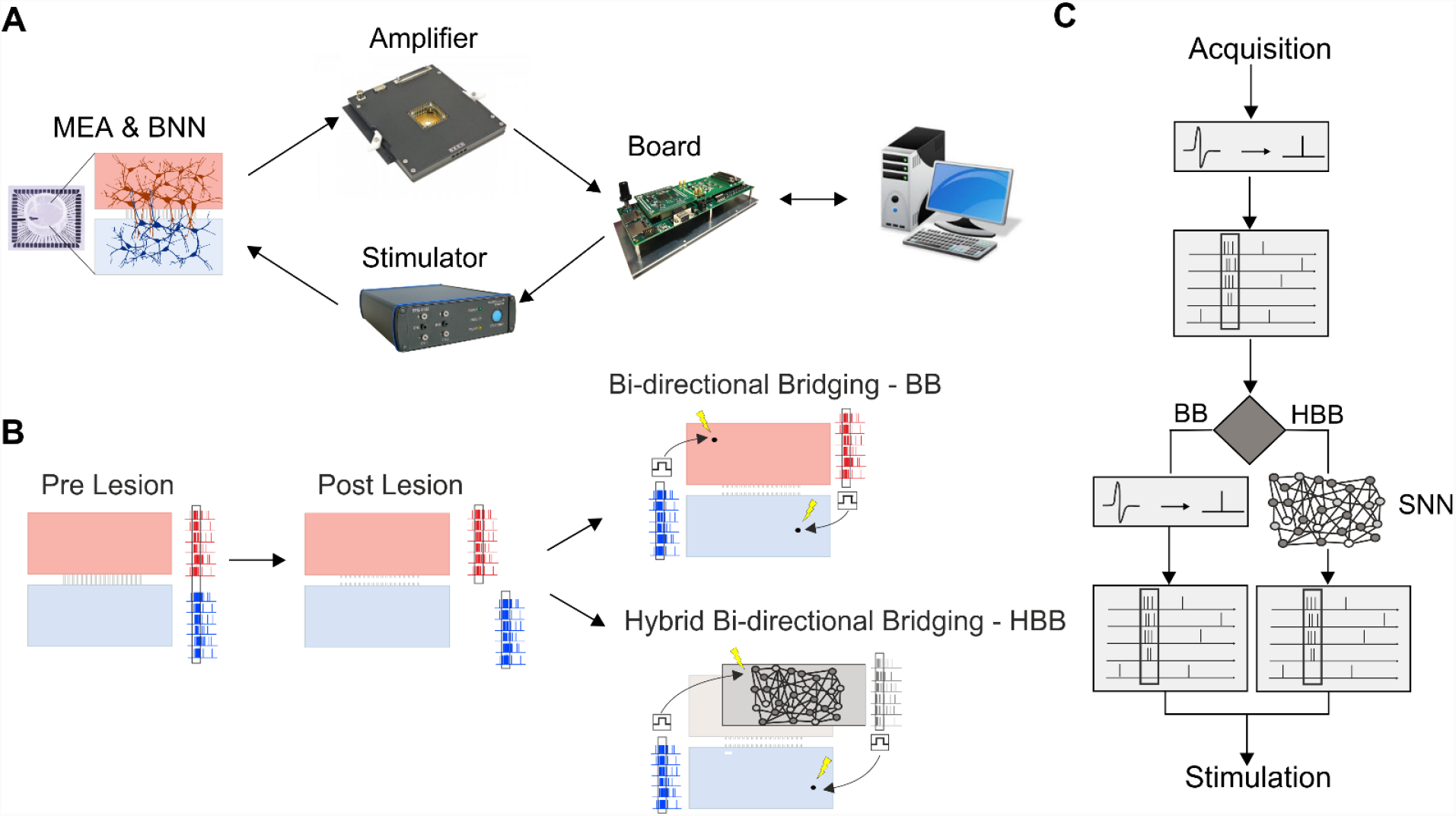
Interfacing a biological neural network and neuromorphic neuroprosthesis. (**A**), Schematic representation of the main elements of the setup: cartoon of an MEA coupled with a bimodular biological neuronal network (BNN); picture of the MEA1060-Inv-BC amplification system (Multichannel Systems, MCS, Reutlingen, Germany); picture of the custom FPGA board; picture of the stimulus generator (STG 4002, MCS). Outside the loop, we used a PC to configure the board. (**B)**, Schematic representation of the different phases of the two experimental approaches used to test our neuroprosthetic device. The first step consisted of checking the healthy condition of the bimodular culture and recording its spontaneous activity (pre lesion). The culture is schematically represented by the top module (light red) connected via light grey segments to the bottom module (light blue). The spiking activity from the two modules is represented by the top red raster plot and by the bottom blue raster plot. Network bursts triggering stimulations are highlighted by a black rectangle. The same color code is used in all figures and panels in the manuscript. The second step consisted of creating a lesion via laser ablation of neuronal connections (ref. video of the laser ablation) and recording spontaneous activity after the lesion (post lesion). The third step was either bidirectional bridging (BB) or hybrid bidirectional bridging (HBB) after the lesion. **c**, Schematic of real-time data processing performed by the board: the first step is spike detection followed by network burst (NB) detection monitoring module 1. After NB detection, delivering stimulation to module 2 of the BNN (BB approach) or to the SNN is possible. In the second modality, there is also NB detection of the SNN, which can result in stimulation delivered to the BNN (HBB approach).

### Control Experiments

To evaluate the stability and effect of the focal lesion on bimodular BNN activity, we first performed two sets of control experiments. In the first set defined as ‘experiments with no lesion’ (Fig. 2A1), we recorded 4 consecutive hours of spontaneous activity (S1-S4, n=9 cultures). In the second set defined as ‘experiments with a lesion’ (Fig. 2A2), we recorded one hour of spontaneous activity (S1), followed by laser ablation of the connections between the two modules, which usually took less than 20 minutes. Next, we recorded 3 hours of spontaneous activity post lesion (SPL1-SPL3) to quantify the effects of laser ablation (n=4 cultures). As depicted in the raster plot of one representative experiment (Fig. 2B1), bimodular neuronal networks exhibited spontaneous, synchronized, multi-unit activity composed of network-wide bursts (NBs) spreading over the two modules. Following laser ablation, the propagation between compartments was disrupted, as shown in Fig. 2B2.

**Fig. 2.**
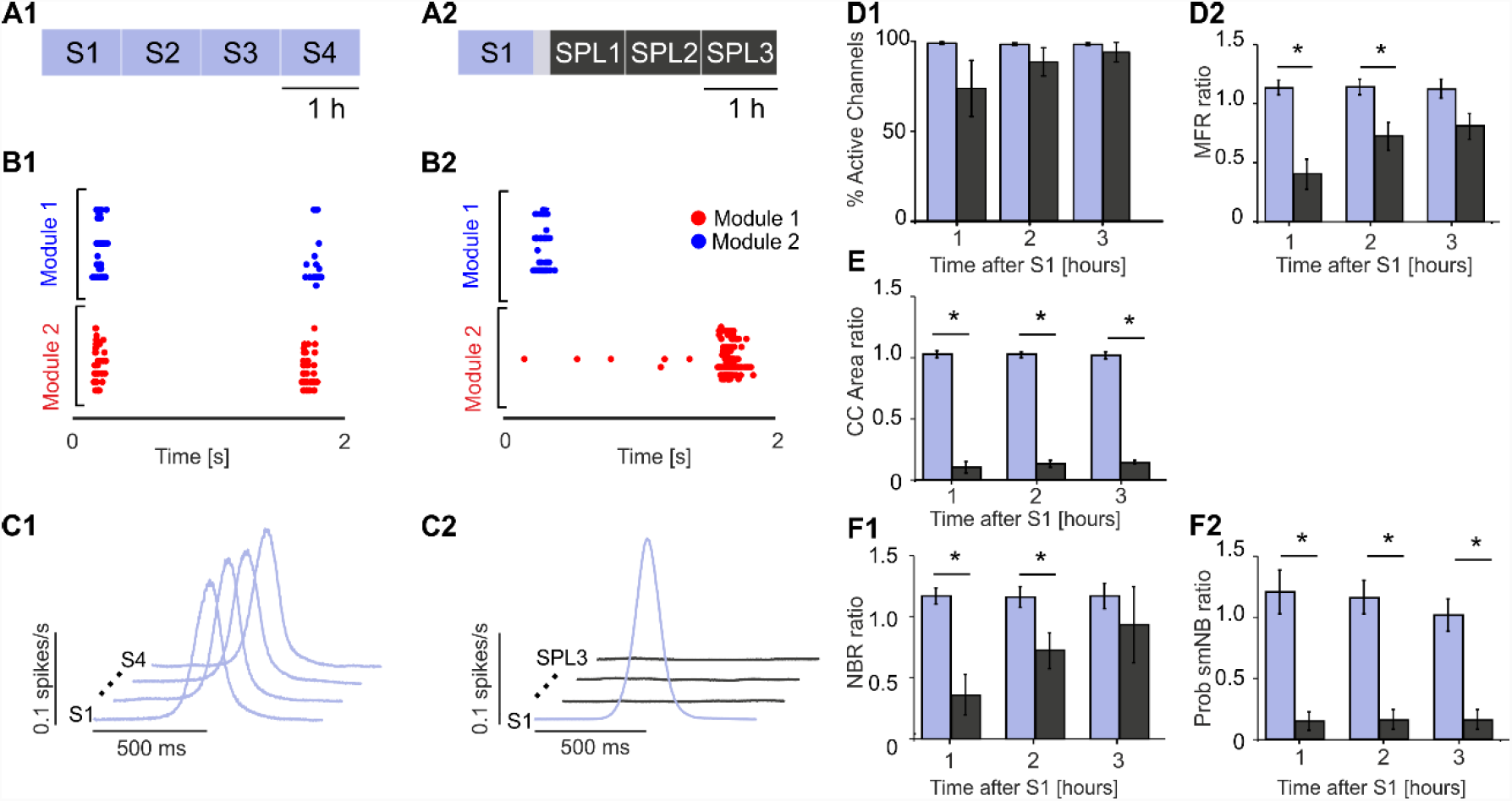
A laser ablation-induced lesion can disconnect two neuronal modules. (**A1**), Schematic of the first experimental protocol. Experiments with no lesion: we recorded four consecutive hours of spontaneous activity (S1-S4). (**A2**), Schematic of the second experimental protocol. Experiments with lesion: we recorded one hour of spontaneous activity (S1) followed by laser ablation and three consecutive hours of spontaneous activity post lesion (SPL1-SPL3). The grey-shaded area indicates 20 minutes of no recording due to the execution of the lesion. (**B1**), A 2 s raster plot of the network bursting activity of one representative experiment during the S1 phase. (**B2**), a 2 s raster plot of the network bursting activity of one representative experiment during SPL3. (**C1**), Cross-correlation (CC) function for one representative experiment during the S1-S4 phases. The CC profiles between the spike trains of each module (light blue) in the four phases of the experiment were high and stable (lines shifted for the sake of clarity). Time axis [-500, +500] ms. (**C2**), CC profile between the spike trains of each module for one representative experiment with lesion. Before the lesion (light blue profile), CC was high; following the lesion (dark grey), CC collapsed to zero (lines shifted for the sake of clarity). Time axis [-500, +500] ms. (**D1**) Percentage of active channels with respect to S1 for the experiments with no lesions (light blue columns, n=9) and with lesions (n=4, dark grey columns). No significant difference was found using the Mann-Whitney test (S2 VS SPL1 p=0.2042; S3 VS SPL2: p=0.31608; S4 VS SPL3: p=0.70769). (**D2**), Mean firing rate (MFR) ratio with respect to S1 for experiments without (light blue bars) and with lesions (dark grey bars). No significant difference was found during the last hour using the Mann-Whitney test (S2 vs SPL1: p=0.0028; S3 vs SPL2: p=0.01119; S4 vs SPL3: p=0.10629). (**E**), Comparison of the CC area ratio with respect to S1 for the experiments without (light blue bars) and with lesions (dark grey bars) (Mann-Whitney test; S2 vs SPL1: p =0.0028; S3 vs SPL2: p=0.0028; S4 vs SPL3: p=0.0028). (**F1**), Network burst rate (NBR) ratio with respect to S1 showing significant differences during the first and second hour after the lesion (Mann-Whitney test; S2 vs SPL1: p=0.0028; S3 vs SPL2: p=0.01119; S4 vs SPL3: p=0.26014). (**F2**), Probability of single-module NB (Prob smNB). The ratio with respect to S1 shows stability for experiments without (light blue bars) and with lesions (dark grey bars) (Mann-Whitney test; S2 vs SPL1: p=0.0028; S3 vs SPL2: p=0.0028; S4 vs SPL3: p=0.0028).

With no lesion, the percentage of active channels with respect to the first hour of recording (S1) was higher than 98% and was maintained for the entire duration of the experiment (Fig. 2D1, light blue bars). The mean firing rate (MFR) was stable for all control experiments with no lesion (from S1 to S4, Fig. S3A). Alternatively, control experiments with lesions showed a reduced number of active channels (close to 73%) during the first hour after ablation (SPL1). During the following two hours (SPL2-SPL3), this value increased and reached 93% at the end of the recording (Fig. 2C1, dark grey bars). No significant differences were found. The MFR was quite stable for all control experiments with lesions except between S1 and SPL1 (Fig. S4A). The activity level with respect to the S1 phase, expressed by the MFR ratio with respect to S1, was stable during the control experiments without lesions (Fig. 2D2, light blue bars). The lesion produced a clear decrease in activity in most cultures, especially during the first two hours (SPL1-SPL2, Fig. 2D2). We found a significant difference between the two experimental sets during the first two hours after S1 but not during the last hour. This result suggests that two hours after a lesion, almost complete spontaneous recovery occurred in terms of the firing rate for the two neuronal modules.

To evaluate changes in the synchronicity between the two modules, we performed cross-correlation (CC) analysis between the collapsed spike trains of each module. The shape of the CC function was stable throughout the entire recordings in experiments without lesions, as reported in Fig. 2C1 for a representative experiment. After a lesion, the CC function collapsed to zero and did not recover during the experiment (Fig. 2C2: representative experiment). To quantify this difference, we integrated the CC function in a range of ±500 ms to obtain the CC area. We did not find any significant change in the CC area values for all experiments with no lesion (Fig. S3B). By contrast, the CC area values showed a marked decrease following the lesion (SPL1). This decrease was due to the lack of anatomical connections between the compartments and did not recover by itself (Fig. S4B). Comparing the CC area ratio between later phases and S1 resulted in a significant difference between matching periods in ‘lesion’ and ‘no lesion’ experiments (Fig. 2E). The network bursting rate (NBR) was stable during all experiments with no lesions (Fig. S3C). For the experiments with lesions, this parameter was less stable but with no significant differences between phases (Fig. S4C). When comparing the two experimental protocols with the NBR ratio with respect to S1, we found significant differences during the first and the second hour post lesion (Fig. 2F1). The mean probability to have NBs composed of spikes belonging to a single module (i.e., Prob smNB, see Methods) was close to 0.2 in the experiments without lesions (Fig. S3D), meaning that the majority of NBs in an intact bimodular network involved both modules. Alternatively, following the lesion, the probability became close to 1 (Fig. S4D), meaning a total loss of functional communication between the two compartments. Using the Prob smNB ratio with respect to S1 (Fig. 2F2), we found significant differences between the two experimental groups during all phases post lesion (Mann-Whitney test; p < 0.05). Thus, for the no lesion experiment, the Prob smNB remained very similar to the initial values, while for the lesion experiments, it changed abruptly due to the lesion. This result further confirmed that the lesion was effective in functionally disconnecting the two modules.

### Experiment 1: Bidirectional Bridging (BB)

The goal of this experiment is to restore communication between two neuronal assemblies after lesion-induced separation. To achieve this goal, we designed and implemented a stimulation reactive paradigm inspired by the ‘activity-dependent stimulation’ (ADS) described in (*21*) in our neuromorphic board. In contrast to the control experiments, the general protocol (Fig. 3A) included 20 minute recordings of spontaneous activity before the lesion (S1). Upon the lesion execution, we waited for two hours to reach stable activity in both modules, as shown by the results of control experiments (cf. Fig. 2). Then, we recorded 20 minutes of spontaneous activity (SPL3). The raster plot of a representative experiment is reported in Fig. 3B. Before the lesion (S1), the bursting activity involved both modules (Fig. 3B1), whereas after the lesion (SPL3), the activity was characterized by single-module NBs (Fig. 3B2). To choose the best parameters (threshold and window time, cf. Methods) that allowed us to reliably detect NBs in both modules, we performed offline NB detection. After the FPGA was updated with these parameters, a 20 minute session of BB was conducted. (cf., Fig. 1B and C for the description of the BB protocol). The BB approach implemented a reactive paradigm; every time an NB was detected in one module, a stimulation pulse was delivered to an electrode in the other module (see Methods) in both directions. During the BB phase, the bursting activity involved both modules similar to the intact condition due to the bidirectional stimulation pulses (Fig. 3B3, blue and red lines represent electrical stimulation pulses delivered from module 1 to module 2 and vice versa). The last phase of the protocol involved 20 additional minutes of spontaneous activity (SPL4), which showed the same activity as SPL3 (Fig. 3B4). We did not observe significant changes in spiking activity (i.e., MFR) throughout the recordings (Fig. 3C).

**Fig. 3.**
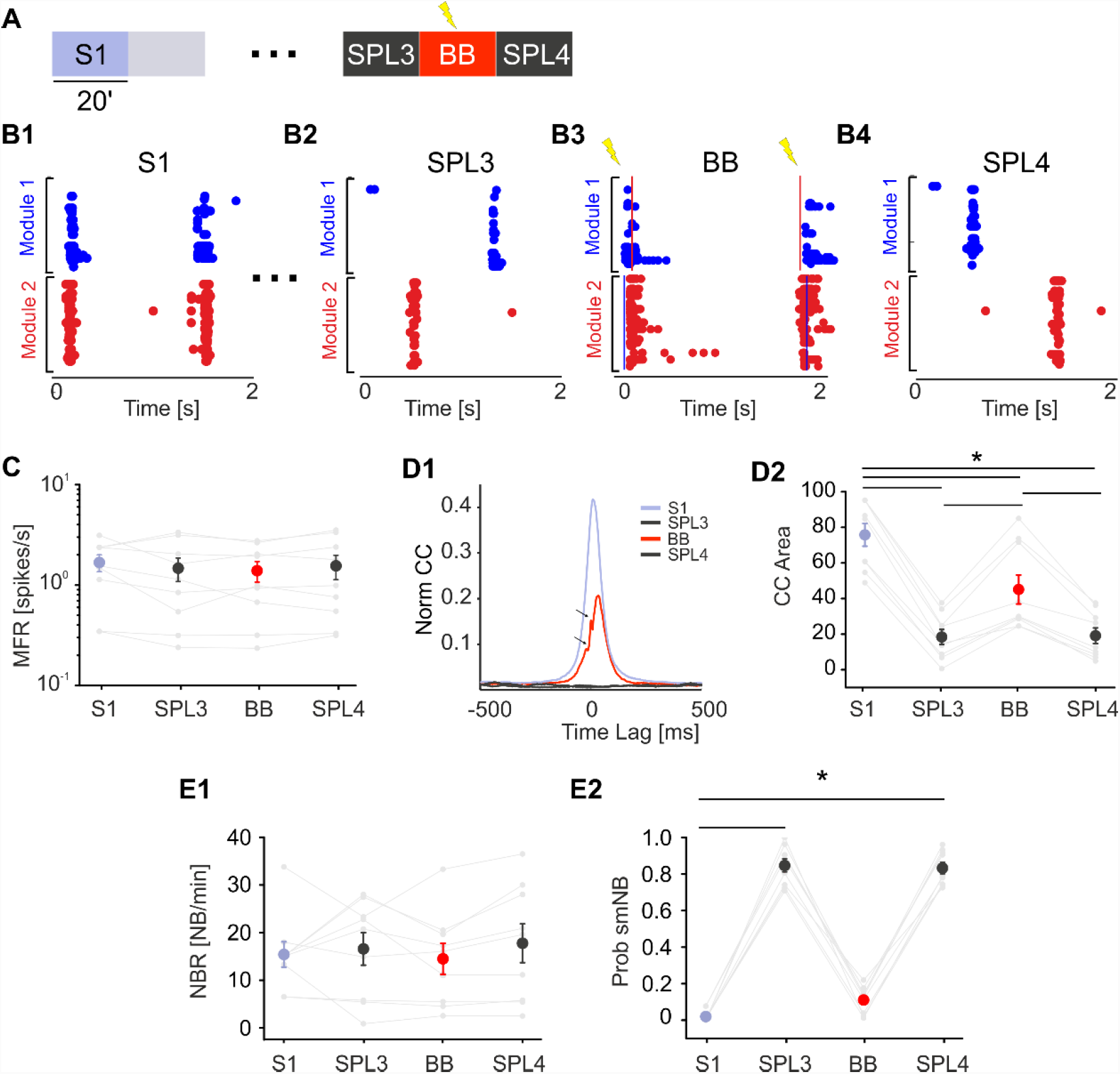
Bidirectional bridging is effective in reconnecting functionally and anatomically disconnected neuronal modules. (**A**), Schematic of the experimental protocol. We recorded 20 minutes of spontaneous activity (S1) followed by laser ablation. The grey-shaded area indicates 20 minutes of no recording during ablation. The dots represent two hours of no recording after the lesion to maintain stable activity in both modules. Then, we recorded 20 minutes of SPL activity (SPL3) followed by 20 minutes of the bidirectional bridging (BB) protocol and another 20 minutes of spontaneous activity (SPL4). (**B1-4**), The 2-s-long raster plots of representative experiments (respectively, from phases S1, SPL1, BB and SPL4). In (**B3**), Blue and red lines represent electrical stimulation pulses delivered from module 1 to module 2 and vice versa, respectively. (**C**), MFR during the 4 experimental phases was stable (color code as in panel a: S1: light blue dot; SPL3, SPL4: dark grey dots; BB: red dot). No significant difference was found (one-way RM ANOVA, p=0.469, DF=3, F=0.872). (**D1**), CC function during the 4 experimental phases. The small arrows indicate the blanking period of 8 ms following each stimulation. The color code is the same as that in panel a. Note that during BB, the cross-correlation function (red) recovers at least partially with respect to the initial profile (light blue), while it stays at zero during the spontaneous activity phases post lesion (SPL3 and SPL4, dark grey profiles). (**D2**), The CC area was highly reduced during the post-lesion phases. The CC area partially recovered during the BB protocol and collapsed again when stimulation was switched off (one-way repeated measures analysis of variance; degrees of freedom=3; F=101,832. S1 vs SPL3 p=5.67E-13; S1 vs SPL4 p=7.54E-13; S1 vs BB p=1.60E-07; BB vs SPL3 p=1.77E-06; BB vs SPL4 p=2.81E-06; SPL4 vs SPL3 p=1). (**E1**), NBR remained stable during the experiments. No significant difference was found (one-way repeated measures analysis of variance: p=0.501, DF=3; F=0.810). (**E2**), The probability of the single-module NB (Prob smNB) was close to one after the lesion. During the BB protocol, the probability was similar to the pre-lesion condition (Friedman’s repeated measures analysis of variance; p<0.001, DF=3, Chi-square=24.3. SPL4 vs S1 and SPL3 vs S1: p<0.001).

Next, we evaluated the effect of this configuration in terms of CC (Fig. 3D1 and 2). During spontaneous activity before the lesion (S1), the CC peak was high and stable due to the functional and anatomical connections between the two modules, which was also reported for the control experiments. After the lesion (SPL3), there was a decrease in CC that was not expected to recover without external intervention, as we previously demonstrated (cf., Fig. 2D). The bidirectional stimulation at least partially recovered the CC area and consequently the communication between modules (Fig. 3D2), as demonstrated by statistical analysis. Regarding the number of NBs, we did not find any significant difference between the experimental phases (Fig. 3E1). However, the probability of isolated NBs was not uniform; it reached the maximum value after the lesion, as we previously observed in the control experiments with lesions (cf., Fig. 2D2). During bidirectional stimulation, these values became closer to the spontaneous recording (Fig. 3E2), meaning that NBs mainly involved both modules. This finding further confirmed that the BB protocol could reconnect two disconnected modules though a real-time ADS acting in both directions (from module 1 to module 2 and from module 2 to module 1).

### Experiment 2: Hybrid Bidirectional Bridging (HBB)

With an injury causing damage to an entire neuronal subnetwork, a reconnection strategy such as the BB illustrated above would not be feasible. For this reason, we developed a second reconnection strategy based on the use of a hardware SNN that can interact in real time with its biological counterpart, HBB (cf. Fig. 1). We created a set of SNNs (i.e., SNN library) by tuning the mean value of the synaptic weight distributions of our models to cover the variability of the BNNs (i.e., BNN library, Fig. 4A). The biomimetic SNN, working in hardware real time to allow bidirectional communication with living neurons, was modelled as a network of 100 Izhikevich (IZH) neurons, with 80 excitatory and 20 inhibitory neurons (cf. Methods (*22*)), according to the biological composition of dissociated cultures (*23, 24*). Synaptic noise (*25*), inhibitory and excitatory synapses (*26*), short-term plasticity (*27*) and axonal delays were included in the model to recreate the network dynamics (cf. Methods and Fig. S6A1 and 2). Regarding the connectivity rules, we set the outdegree (i.e., the number of post-synaptic neurons) to 25 for all neurons in the network, while the indegree (i.e., the number of pre-synaptic neurons) followed a normal distribution with a mean value of 25 and a standard deviation of 4.3 (Fig. S6, R-Square=0.806). The goal of creating an SNN library was to cover a wide range of NBRs because NB was chosen as the triggering event for our reconnection paradigm, as explained above. To this end, we tuned only the mean value of the normal distribution of synaptic weights (the standard deviation was kept constant at the value of 0.3). By increasing or decreasing the mean synaptic weights, we tuned the NBR. For excitatory synapses, the mean value ranged from 0.99 to 1.34 (Fig. 4C left), while that for inhibitory synapses ranged from -2.02 to -1.02 (Fig. 4C right and Table S1). As previously stated, our goal was to cover the NBR variability and not the MFR. The MFR variability in our BNN library was higher than that obtained with our SNN library (Fig. 4E and F1). Nevertheless, the BNN variability in terms of NBR was completely covered by our SNN library, which also contains networks with a much higher NBR than that in the BNN library (Fig. 4F2).

**Fig. 4.**
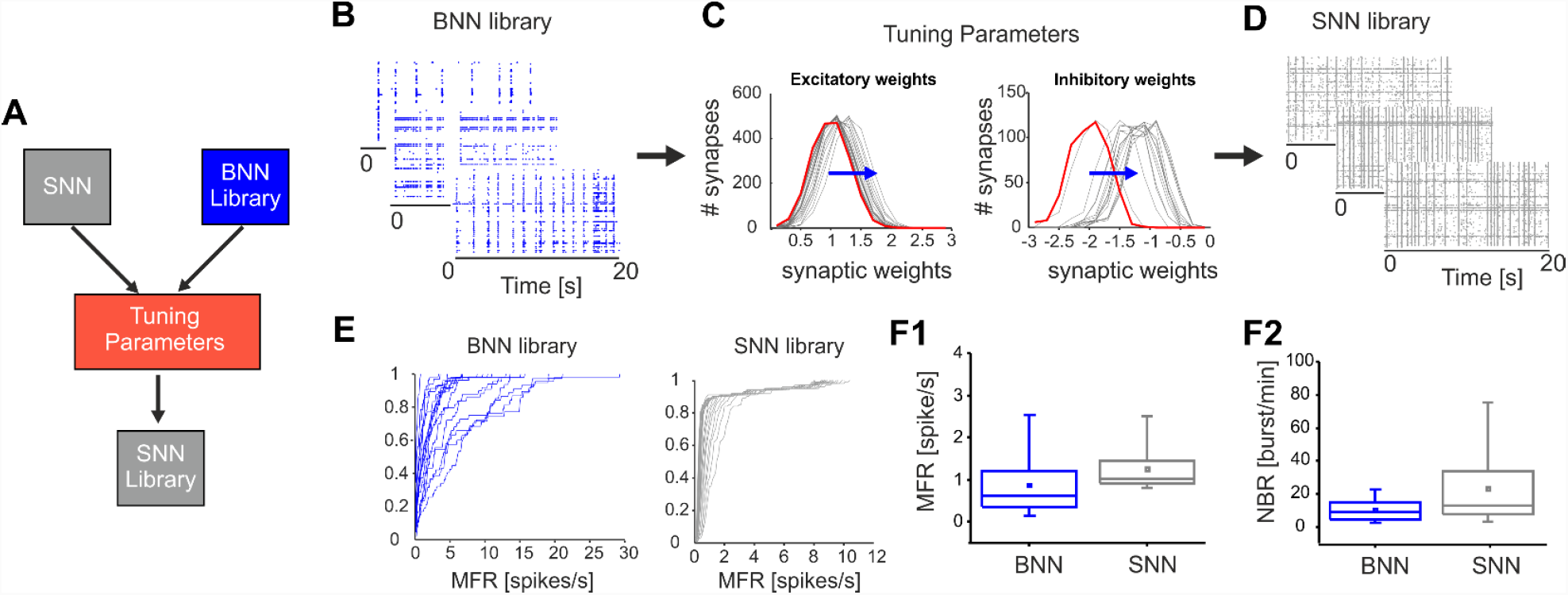
Spiking neural network (SNN) design and characterization. (**A**), Schematic of the procedure used to create a library of SNNs. Starting from the Izhikevich model implemented on the FPGA and a library of 34 BNNs with a large spectrum of activity, we tuned the mean value of the synaptic weight distribution to obtain and select from a collection of SNNs (SNN library, comprising 27 different configurations). (**B**), Representative 20 s-long raster plots of different BNNs showing different NB rates. (**C**), Left, distribution of excitatory synaptic weights from the 27 SNNs. In red, the slower SNN of the library (SNN 1). The blue arrow indicates the shift of the mean value (from 0.99 to 1.34) of the normal distribution with standard deviation = 0.3 to obtain increasing NBR values. Right, distribution of inhibitory synaptic weights from the 27 SNNs. In red, the slower SNN of the library (SNN 1). The blue arrow indicates the shift of the mean value (from -2.02 to -1.02) of the normal distribution with standard deviation = 0.3 to obtain increasing NBR values. (**D**). Representative 20 s-long raster plots of different SNNs showing different NB rates. (**E**), Left, cumulative MFR profile for the BNN library. Right, cumulative MFR profile for the SNN library. (**F1**), Comparison between BNN and SNN libraries in terms of network MFR (i.e., the mean value of all active electrodes for BNN and neurons for SNN). (**F2**), Comparison between BNN and SNN NBR, showing that the SNN library covers the BNN variability and contains networks with a higher NBR.

The general HBB protocol (Fig. 5A) is similar to the BB protocol. The HBB procedure included a 20 minute recording of spontaneous activity before the lesion. This recording was used to quantify activity in terms of the NB rate of the network. This feature was used to choose one SNN from the SNN library that had an NB rate closer to its biological counterpart. We waited two hours after the lesion to allow activity in both modules to stabilize, as shown by the results of control experiments (e.g., Fig. 2). Then, we recorded 20 minutes of spontaneous activity. After setting FPGA detection parameters, we performed a 20 minute HBB session (cf., Fig. 1B and C for the description of the HBB protocol). As anticipated, the HBB approach also implemented an ADS paradigm; every time an NB was detected on the ‘surviving’ module (i.e., when one of the two modules was completely damaged), a stimulation pulse was delivered to the SNN. The board implemented the corresponding paradigm in the opposite direction. Detection of NBs occurred in the SNN, while stimulation was delivered to the ‘surviving’ module, thus avoiding the imposition of any predefined unidirectional communication. Next, we recorded 20 additional minutes of spontaneous activity.

**Fig. 5.**
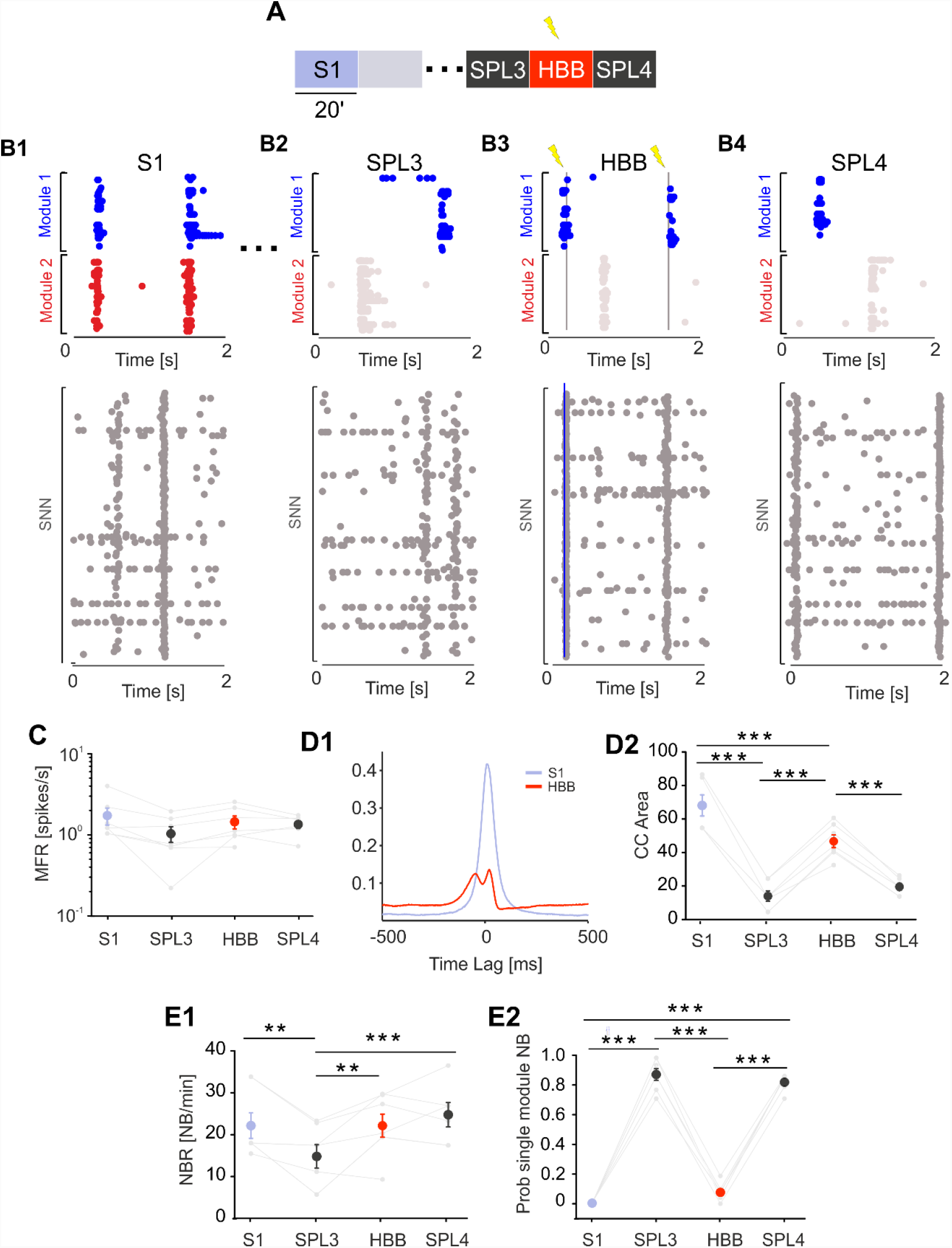
The hybrid bidirectional bridging approach is effective when a neuronal assembly must be replaced. (**A)**, Schematic of the experimental protocols. We recorded 20 minutes of spontaneous activity (S1) followed by laser ablation. The gray-shaded area indicates 20 minutes of no recording during ablation. The dots represent two hours of no recording after the lesion to obtain stable activity in both modules and to test different stimulation channels. Then, we recorded 20 minutes of SPL activity (SPL3) followed by 20 minutes of a hybrid bidirectional bridging (HBB) protocol and another 20 minutes of spontaneous activity (SPL4). (**B1**), Top, 2-s-long raster plot depicting the BNN bursting activity involving both modules before the lesion. Bottom, activity of SNN uncorrelated with the BNN. The networks are not linked. (**B2**), Top, 2-s-long raster plot after the lesion showing uncorrelated bursting activity on BNN modules 1 and 2. Bottom, same as that in b1. (**B3**), 2-s-long raster plot during HBB depicting two hybrid events. The first event on the left was an NB detected on module 1 of the BNN. The detection resulted in a stimulation pulse delivered to 10 excitatory neurons of the SNN (blue line, bottom). An NB on the SNN was detected 18 ms after the stimulation and triggered the delivery of a stimulation pulse to module 1 of the BNN (grey line, top). (**B4**), 2-s-long raster plot depicting the uncorrelated activity of the BNN modules (top) and SNN network (bottom). (**C**), MFR during the 4 experimental phases was stable (color code as in panel a: S1: light blue dot; SPL3, SPL4: dark grey dots; HBB: red dot). No significant difference was found (one-way repeated measures analysis of variance. p<0.001, DF=3, F=3.16; Bonferroni test: all comparisons with p>0.05). (**D1**), CC function during the 4 experimental phases. The color code is the same as that in panel a. Note that during BB, the cross-correlation function (red) recovers at least partially with respect to the initial profile (light blue). (**D2**), The CC area was highly reduced during the post-lesion phases. The CC area partially recovered during the BB protocol and collapsed again when stimulation was switched off (one-way repeated measures analysis of variance. p<0.001, DF=3; F=70,448; S1 vs SPL3: p=5.80E-10; S1 vs SPL4 p=2.72E-09; S1 vs HBB: p=9.16E-04; HBB vs SPL3 p=1.16E-07; HBB vs SPL4 p=1.02E-06; SPL4 vs SPL3 p=0.73643). (**E1**), NBR did not change during HBB with respect to the S1 phase (one-way repeated measures analysis of variance. p=0.005, DF=3; F=6,069; S1 vs SPL3 p=0.02482; S1 vs SPL4 p=1; S1 vs HBB p=1; HBB vs SPL3 p=0.01022; HBB vs SPL4 p=1; SPL4 vs SPL3 p=0.00674). (**E2)**, The probability of a single-module NB (Prob smNB) was close to one after the lesion. During the HBB protocol, the probability was similar to that in the pre-lesion condition (one-way repeated measures analysis of variance. p<0.001, DF=3; F=453,439; S1 vs SPL3 p=4.96E-13; S1 vs SPL4 p=1.03E-12; S1 vs HBB p=0.22606; HBB vs SPL3 p=1.94E-12; HBB vs SPL4 p=4.30E-12; SPL4 vs SPL3 p=1).

We did not observe significant changes in terms of spiking activity (i.e., MFR) throughout the recordings (Fig. 5C). Then, we evaluated the effect of this configuration in terms of CC (Fig. 5D1 and 2). During spontaneous activity before the lesion (S1), the CC peak was high and stable due to the functional and anatomical connections between the two modules, which was also reported for the control experiments. As expected, with no external intervention, CC decreased sharply after the lesion (SPL3), as we previously observed. One of the two modules was damaged, while the correlation was evaluated between the SNN and the surviving module during the HBB phase. The bidirectional stimulation created a relevant correlation area between SNN and the surviving module, as demonstrated by statistical analysis. Regarding the number of NBs, we did not find a significant difference between the S1 and HBB phases (Fig. 5E1). However, the probability of isolated NBs was not uniform; it reached the maximum value after the lesion, as we previously observed in the control experiments with lesions (cf., Fig. 3E2). During the hybrid bidirectional stimulation, these values became closer to the spontaneous recording (Fig. 5E2), meaning that NBs mainly involved both modules. This finding further confirmed that the HBB protocol created a hybrid system with the surviving biological module though real-time ADS acting in both directions (from BNN to SNN and from SNN to the BNN).

## Discussion

We presented an innovative neuroprosthetic system based on a neuromorphic FPGA board and demonstrated two successful reconnection paradigms for a lesion interrupting the communication between two neuronal populations in vitro. This neuromorphic prosthesis and the paradigms implemented herein represent the first example of a successful real-time next-generation neurobiohybrid system (*3*), implementing bidirectional reconnection between two neuronal networks with clear-cut potential for applications to brain injury in humans (*4*), The use of a fully integrated hardware computing system allowed hard real-time performances, which are crucial for translational purposes related to therapeutic applications in humans (*28, 29*).

According to previous reports, in vitro systems constitute a successful experimental model of neuronal dynamics (*30, 31*), thus providing an excellent test bed for adaptive closed-loop neural interfaces (*32*). Starting from our recently developed methodology (*33*), we created custom bimodular cultures with the goal of reproducing two interacting neuronal populations, thus mimicking the intrinsic modularity of the brain (*17, 34*). Our bimodular cultures were highly temporally stable in terms of firing properties at the whole network level as the activity between the two populations remained highly correlated for the entire duration of the recording. A lesion produced via laser ablation was employed to physically cut the connections between the two modules. As previously demonstrated, this methodology is safe because it produces localized damage by selectively ablating subcellular compartments without damaging adjacent structures (*18, 35, 36*). This technique proved successful in decoupling activity in the two modules while preserving their functionality, as demonstrated by the complete recovery of firing rates and non-existent correlation of bursting behavior.

Two different applications of our neuroprosthesis, BB and HBB, were tested. Our neuromorphic prosthesis, independently of the stimulation paradigm, works according to a closed-loop reactive policy as follows: each time a condition is met (i.e., an ‘event’ is detected), a stimulus is delivered. The proof-of-principle provided in this work allows direct application to more complex animal-based models (such as ex vivo brain preparations and in vivo) and in the future, in humans as neuroprosthetic devices for treating brain lesions. To this end, the hardware architecture was designed to be flexible enough to allow the implementation of different experimental paradigms and the definition of different triggering events. In our study, we chose ‘NBs’ as trigger events (cf. Methods). The choice to deliver a stimulation depending on a network-wide event has two main advantages as follows: first, NB frequency is low enough to avoid inducing plasticity phenomena by electrical stimulation in our cultures (*37*), which could confound the final results and effectiveness of neuroprosthetic reconnection. The second point is anticipation of the following major issue that will emerge during in vivo recordings: monitoring single neurons presents problems at both theoretical (*21*) and practical levels. Namely, how much information on complex functions can be obtained by single-neuron observation remains unclear (*38, 39*), while tracking the activity of the same neuron for extended periods of time is problematic (*40*). Taking multiple input sources into account was also used in the work of Berger et al. (*41*), but they employed a neuroprosthetic strategy different from ours. In particular, these authors used a generalized linear model to predict the CA1 activity from spikes in CA3 of the hippocampal circuit. Our system is considered more flexible and adaptable to networks with different connectivity, not just feedforward similar to that in the hippocampus. Moreover, we were interested in mimicking the overall spiking activity of the network and not mapping an input-output transformation only.

Another important novelty of our system regards directionality. To our best knowledge, this neuroprosthetic system is the first to implement a truly bidirectional interaction through a hard real-time interface. We recorded activity from the first module (via multiple sources); when a criterion was met, the device stimulated the second module (this is how a ‘typical’ closed-loop in neuroscience works, for a review see (*42*)). The novelty is simultaneously monitoring multiple sources from another module and delivering the stimulation when the triggering event is detected. To date, all neuroprosthetic devices that have been proposed in the literature can implement a ‘unidirectional’ artificial link only from one area to another (or maybe the same) but not doubling it. Here, we are not imposing any preferred directionality to the communication; networks are self-organizing on the basis of their intrinsic natural relationship (we are not imposing who is driving whom). This approach has the main advantage of informing both brain regions (i.e., neuronal modules, in our case) that an event occurred in the other region, given that interaction in the brain is intrinsically bidirectional (*43*). For example, in the sensorimotor system, sensory simulation can help motor recovery (*44*), and motor learning can enhance sensory functions (*45, 46*). Applications of our neuroprosthetic systems to conditions where the sensorimotor interaction is impaired would allow restoration of both communication channels, suggesting improvements in current rehabilitation therapies. Moreover, although tested on a bimodular system, the neuromorphic FPGA board can be easily upgraded to play the bridging role on an arbitrary number (within reason) of different neuronal circuits.

The second paradigm we tested was based on the use of a biomimetic SNN to ‘substitute’ a missing/damaged neuronal population. Currently, SNN applications span different fields, including computational neuroscience (*47, 48*), and very recently, they were used for sensory encoding in hand prosthesis for amputees (*49, 50*). SNNs can be simulated in software (*51*) and/or neuromorphic hardware. As time and energy consumption are fundamental in neuroprosthetic applications for translational purposes, the use of hardware-based computing systems becomes mandatory.

In general, hybrid systems composed of in vitro BNNs coupled to SNNs are rare. In one approach, the SNN served as a self-organizing classifier of activity patterns exhibited by the BNN, with output of the SNN being subsequently used to control the behavior of a robot (*52*). Other studies focused on the unidirectional or bidirectional influence of the two networks, investigating the dynamics of the interaction between the BNN and SNN in which the SNN played a role of an artificial counterpart of its biological original (*53, 54*). However, closed-loop effects in those hybrid networks were not thoroughly determined. In one of these studies, only unidirectional connectivity was considered with input from the SNN to the BNN, which was also simulated beforehand (*53*). In this study, we established hybrid communication in the case of an entire neuronal population that needed substitution.

A study by Chou et al. (*54*) implemented a bidirectional interface between an SNN and a retinal slice obtained from an adult rat and recorded by an MEA. This system is quite interesting, but there is a 1 s delay between the BNN and SNN interactions. Therefore, this delay in Chou’s setup is 3 orders of magnitude larger than that in our work in which the sampling of biological activity is never interrupted, and the step size of the SNN is 1 ms. The difference between the two systems is radical; bidirectional communication in real-time allows actual clinical application, whereas delays in the range of seconds prevent (or at least seriously reduce) the possibility of meaningful control of a biological system.

A recent study, inspired by a previous work (*55*), implemented a hybrid interaction (*56*) between the cerebellum of a rat and an SNN implemented on FPGA. Their model involved 10k LIF neurons and did not integrate other biomimetic behaviors, such as axonal delay, short-term plasticity and synaptic noise, unlike the IZH neurons implemented in our system. Both the hard real-time processing and simplified neuronal model (which allow mimicking the richness of the electrophysiological patterns in vivo) are mandatory for reproducing the biological dynamics of living neural networks and for performing useful real-time hybrid experiments.

A limitation of the present work is that we deliberately chose to downsample both the number of biological neurons recorded through a low density MEA and the number of artificial neurons implemented on the FPGA. For the purpose of detecting network-wide activity, this oversimplification of biological complexity can be acceptable to test the functionality of the device and the feasibility of the approach, but if the goal is to functionally replace a biological network, a higher resolution would be preferable. Another limitation regarding the SNN is that our implementation lacks a long-term memory function. This aspect can be especially relevant for long-term experiments.

In conclusion, we have proven the possibility of using bimodular cell cultures as a test bench for the development of novel neuroprosthetic systems for therapeutic purposes. We evaluated the loss in communication between the modules following a focal lesion. We recovered the communication between the modules through a custom-developed neuromorphic neuroprosthesis based on an FPGA board, which achieved hard real-time performances. We suggested two promising translational paradigms to improve the current drawbacks of current closed-loop neural interfaces, and we successfully tested them on a biological model of interacting neuronal subpopulations. Our neuroprosthetic system implements a biomimetic SNN exhibiting the same NB dynamics of its biological counterpart. Our experiments demonstrated that the developed neuromorphic neuroprosthesis could restore bidirectional communication between two previously connected neuronal assemblies, thus introducing new applications for brain injuries.

## Materials and Methods

### Bimodular Confinement

The polymeric structure for the physical confinement of neuronal cultures on conventional MEAs has been realized in polydimethylsiloxane (PDMS) by soft lithography and by using a photolithographically defined EPON SU-8 master on a silicon substrate. PDMS is an elastomer widely used for biomedical applications because of its high temperature, chemical and oxidation resistance; biocompatibility; transparency; and permeability to gases (*57*). Additionally, PDMS is an electrical insulator and nontoxic. These features also render PDMS particularly suited to the culture of primary neurons (*58*). In addition, PDMS can be easily micro-structured by soft lithography, thus obtaining several low-cost replicas from a single master (*35, 59, 60*). In our case, replicas were produced for single use even if they could in principle be sterilized and re-used. In particular, PDMS stencil fabrication consisted of the following consecutive steps: i) structure design; ii) in silico wafer fabrication through a photolithographic technique; and iii) fabrication of PDMS replicas (for details, see (*15*)). The developed modular neuronal networks were composed of two 3x2 mm modules that were interconnected by 25 channels 390 µm long and 10 µm wide with an inter-channel distance of 50 µm. Once the PDMS masks were ready, they were positioned on the MEA substrates to include all the electrodes within the two modules. Next, the MEAs underwent the coating procedure for promoting cell adhesion, and the following day, the PDMS mask was removed immediately before the plating procedure, thus allowing neurons to slowly move only towards the promoting adhesion areas.

### Neuronal Preparation

Dissociated neuronal cultures were prepared from the neocortex of 18-day-old embryonic rats (pregnant Sprague-Dawley female rats were obtained from Charles River Laboratories). All experimental procedures and animal care were conducted in conformity with institutional guidelines in accordance with the European legislation (European Communities Directive of 24 November 1986, 86/609/EEC) and with the NIH Guide for the Care and Use of Laboratory Animals. For our experiments, we used dissociated neurons arranged on a bimodular layout (cf. ‘Bimodular confinement’ paragraph for details) grown on MEAs. Culture preparation was performed as previously described (*15, 61*). First, we coated MEAs overnight with poly-D-lysine and laminin to promote cell adhesion (Fig. S1 and A1). We washed the MEA devices at least 3 times with sterilized water before plating (Fig. S1 and A2). The neocortex of 4–5 embryos was dissected from the brain and dissociated first by enzymatic digestion in trypsin solution 0.125% (25–30 minutes at 37°C) and subsequently by mechanical dissociation with a fire-polished pipette. The resulting tissue was resuspended in Neurobasal medium supplemented with 2% B-27, 1% Glutamax-I, 1% Pen-Strep solution and 10% Fetal Bovine Serum (Invitrogen, Carlsbad, CA) at a final concentration of 500 cells/ul (Fig. S1 and A3). Cells were then plated onto 60-channel MEAs and maintained with 1 ml nutrient medium (i.e., serum-free Neurobasal medium supplemented with B27 and Glutamax-I). Then, cells were placed in a humidified incubator with an atmosphere of 5% CO_2_−95% air at 37°C. Half of the medium was changed weekly.

### Focal Lesion Procedure and Laser Setup

The laser dissection system was previously described (*62*). The developed system allowed for focal ablation of the sample in a three-dimensional confined volume due to the sub-nanosecond pulsed UVA laser source, which required delivering very low average power, thus confining the material breakdown to the focus spot. The setup was configured in an upright optical layout to allow optical surgery of neuronal networks plated on thick and non-transparent support such as the MEA chip. Therefore, simultaneously monitoring network activity using the MEA device and fluorescence calcium imaging during optical dissection of the connection of neuronal assembly was possible. The system was equipped with a custom micro-incubator (*63*) maintaining the physiological parameters of the neuronal cultures (pH, osmolarity and temperature) to carry out long-term network activity recording before and after laser injury (*36*).

### Micro-Electrode Array Recordings

MEAs (Multi Channel Systems, MCS, Reutlingen, Germany) consisted of 60 TiN/SiN planar round electrodes (electrode diameter: 30 μm; inter-electrode distance: 200–500 μm). One recording electrode was replaced by a larger ground electrode. Each electrode provided information on the activity of the neural network in its immediate area. The array design using our bimodular mask allowed us to confine the growth of cells in two macro areas hosting 28 electrodes each (i.e., the ground electrode and the three central electrodes were excluded). A microwire connected each microelectrode of the MEA to a different channel of a dedicated amplifying system with a gain of 1100. The signals from the BNN were amplified by a commercial system (MEA1060-Inv-BC amplification system, Multi Channel Systems, MCS, Reutlingen, Germany).

To reduce the thermal stress of the cells during the experiment, we maintained MEAs at 37°C with a controlled thermostat (MCS) and covered them with a custom PDMS cap to avoid evaporation and prevent changes in osmolarity (*64*). Additionally, we used a custom chamber to maintain a controlled atmosphere (i.e., gas flow of 5% CO2 and 95% O2 + N2) during the entire recording time (*65*).

### Electrical Stimulation

All electrical stimuli to BNN were triggered by our FPGA-based board (by a commercial stimulator, STG 4002, Multi Channel Systems, MCS, Reutlingen, Germany). The basic electrical stimulus was a biphasic voltage pulse, 300 µs in the half-length phase and 750 mV of half-amplitude (in accordance with the literature (*66*)). The necessary condition to choose the stimulating electrode was its capability to evoke a clear response on that module, meaning a burst involving the entire module. In the pre-lesion condition, the stimulation evoked a response in both modules. In the post-lesion condition, the stimulation evoked a NB confined to a single module.

### Experimental Protocols and Databases

The general protocol included at least 30 minutes of no recording immediately after the culture was moved from the incubator to the amplifier to ensure the stability of the network in recording conditions. For control experiments without lesions, the four hours were recorded continuously. In controls with lesions, after the first hour of recording, we took the culture from the amplifier system, and we moved it to a separate room for laser ablation. The entire procedure usually took 20 minutes. After the lesion, the culture was moved back to the amplifier system where we immediately started the recording. The total number of experiments performed with these protocols was 9 for controls without lesions (26.8 ± 0.8 days in vitro, DIV) and 4 for controls with lesions (26 ± 0.6 DIV). For the BB protocol and HBB, we performed similar protocols. We recorded 20 minutes of spontaneous activity before the lesion. Next, we performed laser ablation as described above and then waited for two hours to achieve stable activity in both modules, as shown by the results of control experiments (Fig. 2). Then, we recorded 20 minutes of spontaneous activity after the lesion. To choose the best parameters that allowed us to reliably detect NBs in both modules, we performed offline NB detection. Next, we set these detection parameters on the FPGA with a custom-made MATLAB code and performed a 20 minute session of BB or HBB. The final step consisted of recording 20 minutes of spontaneous activity. The total number of experiments performed with these protocols was 9 for BB (26 ± 0.8 Days in vitro, DIV) and 7 for HBB (26 ± 1.5 DIV). In one HBB experiment, we did not record the last spontaneous phase because of a technical problem.

### Neuroprosthetic System and Online Processing

The custom experimental system retrieved analogue data from MEA1060-Inv-BC pre-amplifiers through a standard MCS connector. Analogue input signals were packed in subgroups of 8 signals. For each subgroup, the signals were connected to the input of an analogue multiplexer that switched from one signal to the next one at a frequency of 80 kHz (each 12.5 µs) as shown in Fig. S2A. Then, the output signals were amplified. The amplification gain ranged from 1 (no amplification) to 100. Multiplexed signals were finally converted to digital with 16-bit accuracy at a frequency of 80 kHz to provide one sample each time the multiplexer switched to a new input. Then, each of the 8 inputs of each multiplexer was sampled at a frequency of 10 kHz. The system relied on the antialiasing filters embedded in the pre-amplifier.

The spike detection module of the neuroprosthetic device was based on a threshold principle. A detection system was available for each input channel of the system. The first step of the detection was to increase the signal to noise ratio (SNR) of the input signal. The second step was to determine the appropriate threshold to perform the detection; this part of the processing is illustrated in Fig. S2B. To emphasize the amplitude of spike shapes and improve SNR, we used the wavelet decomposition technique (SWT) as previously described in (*67*). This technique consisted of applying two orthogonal filters (one highpass and one lowpass) tailored to the characteristic shape of a spike. A more thorough description is provided in the supplementary files.

Because of the impedance variation phenomenon, the input gain was expected to be different from one electrode to another and from one experiment to another. Therefore, we needed automatically set thresholds to avoid setting this parameter for each input channel individually. We determined these thresholds as a multiple of the standard deviation (σ) of the first detail level from the SWT outputs. The module that computed σ was referenced as an amplitude estimator (AE) in Fig. S2. The correspondence factor to determine the threshold was set system-wide by the experimenter depending on the experimental conditions. To compute σ, we used a continuous evaluation to avoid characteristic steps that occurred when performing computation on windows. To reduce computational resources, we digitized a method from analogue computing (*68*). With the hypothesis that the signal distribution was constant, we knew the proportion of samples below and above the standard deviation, and the system used this property to determine σ with a regulation loop. The signal distribution may change from experiment to experiment, but the estimated σ will always be relevant to the signal amplitude. Then, variations are compensated by adjusting the multiplier that determines the final threshold. Harrison (*68*) showed that in addition to its computation efficiency, this method was much more immune to accidental phenomena such as spike residues or stimulation artefacts.

As previously stated, the threshold was applied to the chosen output detail level of the SWT. The final step was to filter glitches that may come from noise around the threshold value and to change a spike-length pulse to a single event. This step was performed by a state machine that as far as the threshold had been crossed, provided a detected spike event and remained silent during a user-configured refractory period. Then, the output event was available as a spike event for the remaining computation modules of the system.

### Real-Time Network Burst Detector (NBD)

The burst detector was based on the number of action potentials. Although this module was usable on a single channel, it was intended to gather events from multiple channels at the network scale as the preliminary experiments raised the necessity. Therefore, this module was not part of the computation flow build for each channel. There were 16 NBDs. Each NBD received events produced by all the incoming channels. Whether the input channels contributed to any of the 16 NBDs was user defined. During a predefined time window, events on the selected channels were counted. At the end of the time window, the number of events was compared to a user-defined threshold to determine if an activity burst was occurring or not, and the counter was reset. The detector constantly kept track whether a burst was pending or not. Depending on user choice, each NBD could be independently configured to produce a single event when a burst started, a single event when a burst stopped, one event at each end of computing window detecting a burst, or continuously while the detector was in burst mode.

An offline version of the NBD module was written in MATLAB to expedite the choice of parameters for reliable NB detection during experiments. By running the offline NBD with a combination of different windows in the range 1-50 ms and different thresholds in the range 1-250, we looked for a combination of parameters that could reliably detect NBs.

### Spiking Neural Network

The biomimetic SNN is a neuromorphic system with a most detailed level of analogy with the nervous system. The SNN is a network of silicon neurons connected via excitatory and inhibitory silicon synapses and plasticity rules. The characteristics of the systems for bio-hybrid experiments are biological real-time, complex neuronal models and plasticity that reproduce spatio-temporal patterns of activation. The hardware real-time implementation needs a low-resource neuron model.

### Neuron Model

The (*22*) IZH model is composed of two equations in which ‘v’ is the state variable representing the membrane potential of a neuron and ‘u’ represents the regeneration of the membrane potential, which takes into account the activation of ionic currents K^+^ and the inactivation of ionic currents Na+. The IZH model can reproduce the behavior of several neurons by changing 4 parameters *a, b, c* and *d* (Fig. S6). The computation time of the IZH model is 1 ms. Using a calculation pipeline (Fig. S5B1), we can compute different neurons in parallel. All model parameters are stored in Random Access Memory (RAM). The detailed model and its implementation in digital hardware (*69*) are described in the supplementary files.

### Synapse

Synaptic transmission includes both excitatory and inhibitory models. In particular, the proposed model takes into account AMPA (i.e., an excitatory neurotransmitter) and GABA (i.e., an inhibitory neurotransmitter). Depolarization or hyperpolarization are represented by a positive or negative contribution to synaptic currents. Following the effects of AMPA and GABA, all excitatory or inhibitory synaptic currents tend to zero out, exponentially decreasing (*70*). These two synaptic currents obey the same law of exponential decay (3 ms for excitatory decay and 10 ms for inhibitory decay). According to the literature (*26*), whenever a pre-synaptic neuron emits a peak, a synaptic weight (Wsyn) is added to the synaptic current of the post-synaptic neuron. To improve the biological behavior of our synapses, we added short-term synaptic plasticity, which modified the synaptic weight according to the activity of pre-synaptic neurons.

### Short-Term Synaptic Plasticity Model

Short-term plasticity is a biological phenomenon that modifies and regulates connection weights as a function of network activity. The model used in this work was proposed by (*27*). This model is called “short-term plasticity” because the facilitation as well as the depression of a synapse is reabsorbed when no action potential has been emitted during a time constant. The implementation of the short-term plasticity and synapses are described in the supplementary files (parameters: xsyn, P and tsyn in Supplementary Information).

### Synaptic Noise and Axonal Delay

To enable spontaneous activity and the activity of our network in a more neuro-realistic system, we added stochasticity by a source of current noise in the neuron model. We used the Ornstein-Uhlenbeck (OU) process (with parameters: mean value = 0; degree of volatility = 35; dissipation rate = 1), which is a suitable model for modelling synaptic noise in a neural network (*71*). The OU process Xt is a prototype of a noisy relaxation process and an example of a Gaussian process that has bounded variance and a stationary probability distribution. The process is stationary, Gaussian and Markovian. The digital implementation of this OU process is described in (*25*). Fig. S5C represents the dynamic behavior of the neuron model (tonic bursting neuron) family in which the parameters are set for tonic bursting activity. This figure represents the same neuron in which we add the *stochastic* input currents. We noticed that the current noise source caused variability in the timing of action potentials.

The synapse computation core also manages the axonal delay. The axonal delay is the phenomenon according to which an action potential emitted at time t arrives at time t + t1 to the post-synaptic neuron. In our implementation, the axonal delay was implemented by a 50-bit delay (shift register) vector, a D value and a multiplication factor. We stored the action potential (1 bit) in this vector delay at the position indicated by the value D. We can have a delay from 0 to 49 ms if the multiplication factor is equal to 1. We can increase this time by tuning this factor. The axonal delay makes it possible to describe a superposition of a network of neurons in 2D and consequently design pyramidal or 3D networks.

### Neural Network Architecture

Our architecture of the neural network was based on blocks of RAM (storage of the parameters necessary for the definition of a network), two computing cores (a neuron and synapse), a block to manage the state machine and addresses and an “RS232 Communication” block (which will allow us to configure/modify the parameters of our system from a configuration file). Fig. S5a shows the interaction between these blocks. To simplify the figure, we did not represent the clock signal, but the entire architecture is synchronous. The limitation of the number of neurons is due to the following factors: 1) the clock frequency used to maximize the number of neurons in parallel and 2) the size of the RAM, which allows the storage of the parameters, especially the synapses. The resources used are presented in Table S2.

One of the advantages of our architecture was that each neuron and synapse were independent. A network configuration file was sent to the FPGA to select the connections and set the number of neurons, synapses, and used options. We could also define multiple networks with different connectivity between networks and within each network. This process made it possible to define ’3D’ layer networks similar to those found in biology (pyramidal structures).

### Network Configuration

To add flexibility to our system, we added the ability to communicate with the FPGA via serial links to send the configuration, topology, and neural network options. Several options were possible depending on the desired application as follows: axonal delay, short-term plasticity, and synaptic noise. Then, we chose the family of neurons (excitatory, inhibitory), the percentage of connectivity between neurons, the inhibitory/excitatory ratio k, synaptic weight distributions, and the amplitude of the OU noise. From a library of configuration files, we could select the desired network for the given application and particularly in terms of frequency of NB. Fig. 4 describes SNN library processing. Table S1 shows some metrics relative to the bursting features of the implemented artificial networks as well as the mean values of the excitatory and inhibitory weights.

Depending on the FPGA board, the maximum number of neurons varied. For instance, in a Spartan 6 FPGA board, we implemented 512 neurons, 66048 synapses with synaptic noise, axonal delay and synaptic plasticity. Table S2 describes the resources used. This biomimetic digital SNN worked in real time, was tunable and reproduced complex biological neural networks due to the biophysically neuron model, synapses, plasticity, axonal delay and synaptic noise.

### Offline Data Analysis

Offline data analysis was performed by custom scripts developed in MATLAB (MathWorks, Natick, MA, USA).

### Preprocessing

To characterize the activity level of BNNs, we used the percentage of active channels and the MFR. To compute the former, we considered active electrodes only those presenting a firing rate higher than 0.01 spikes per second (spikes/s), while the latter was defined as the mean number of spikes per second, computed over the total recording time, of the active channels. The low threshold guaranteed excluding only those electrodes that were not covered by cells or with very few spikes, keeping all the others (*15*). In stimulation phases, the activity of stimulated channels was deleted.

### Cross-Correlation

To quantify the level of synchronization among multi-unit recordings, we applied correlation analysis (*72*) to the spike trains. Cross-correlograms were built according to the method of activity pairs described by (*73*) as follows: given two trains (i.e., X and Y), we counted the number of events in the Y train within a time frame around the X event of ±T (T set at 500 ms) using bins of amplitude Δt (set at 1 ms). The CC function Cxy(τ) was obtained by a normalization procedure according to the following formula (*74*):

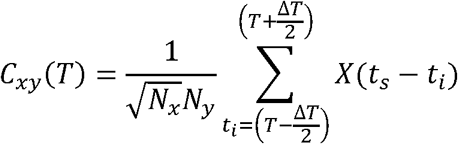

where ts indicates the timing of an event in the X train, Nx and Ny are the total number of events in the X and Y trains, respectively, and Δτ is the bin size. Equation (1) yields the symmetry between Cxy(τ) and Cyx(τ) (i.e., Cxy(τ)=Cyx(-τ)) (*73*).

To evaluate the amount of correlation between two modules, we performed the computation between the collapsed spike trains for each module. A collapsed spike train is the result of a logic OR operation between the spike trains recorded on all the electrodes belonging to the same module. For control experiments and BB experiments, the modules were modules of the BNN. In the HBB protocol, we collapsed the activity of the entire SNN to a single spike train. We applied a locally weighted linear regression to smooth the curve. The area of the CC was the integral of the curve in the range from -500 to 500 ms.

### Offline Network Burst Detection

The offline NBD was performed using a software implementation of the NBD algorithm running online on the neuromorphic device, called the “fixed window accumulator”. Similar to the online version, the algorithm summed all the spikes fired from the neurons of the networks during a given time interval, and it detected an NB if this sum overcame a threshold. Each NB was highlighted by the generation of an event. The input parameters were the window time (i.e., time interval within which the spikes were summed), the threshold (i.e., the number of spikes to overcome to identify an NB event) and the source (i.e., the channels on which the analysis was performed).

Depending on the specific protocol we wanted to analyze, the source could be the whole BNN (in controls without lesions or pre-lesion spontaneous activity), a part of it (for example, to compute the NBR in only one module or to exclude stimulated electrodes), the SNN (for example, to characterize the SNN) or a hybrid network formed by both the SNN and whole BNN or part of it (in HBB). The offline version was also useful during the experimental protocols to set the best parameters (window time and threshold) to obtain a reliable NB detection and to choose the most suitable SNN, referring to NBR, for the HBB experiment.

### Probability Single-Module Network Burst

To quantify the confinement of the electrophysiological activity in a single module, we used the single-module NB probability parameter. This parameter represented the probability that NBs were composed of spikes belonging to a single module (i.e., one of the modules of the BNN or the SNN). This parameter was computed by counting the number of spikes that belonged to each burst, followed by the percentage of spikes that belonged to each module. If this percentage was higher than 85%, we assumed that the burst was generated in a single module. Setting 85% as the threshold percentage allowed us to consider the possibility of having random spikes in the other module at the same time as the NB. To perform this analysis, we modified the NBD by adding start and stop thresholds. For each NB identified, the algorithm looked for an empty time window (i.e., a time window in which the number of spikes was equal or lower than the threshold) before and after the window in which an NB was detected to identify the first and last spikes of that event as the first spike in the first window and the last spike in the last window belonging to the NB, respectively. We set the start and stop thresholds to zero and 5, respectively, for BNN and SNN (the SNN threshold was higher because of the presence of fast spiking neurons).

### Statistics

Data within the text are expressed as the mean ± standard error of the mean (SEM), unless otherwise specified. Statistical tests were employed to assess significant differences among different experimental conditions. The normal distribution of experimental data was tested using the Shapiro-Wilk normality test. We performed one-way repeated measures ANOVA to compare data from the same group at different time points. When ANOVA gave a significant (p < 0.05) result, the post hoc Bonferroni test was employed to assess differences between all phases. When normality was rejected, we used Friedman’s repeated measures ANOVA on the ranks test to compare data from the same group at different time points. When ANOVA gave a significant (p < 0.05) result, the post hoc Tukey test was employed to assess differences between all phases. To compare data from two different populations (Fig. 2), we performed the Mann-Whitney test. Statistical analysis was carried out by using OriginPro (OriginLab Corporation, Northampton, MA, USA) and Sigma Stat (Systat Software Inc., San Jose, CA, USA).

## Acknowledgments

The authors would like to thank Dr. Daisuke Ito, Dr. Marina Nanni, Dr. Claudia Chiabrera, and Dr. Giacomo Pruzzo for precious technical support in performing the in vitro experiments at IIT. The authors are grateful to Dr. Marianna Semprini for useful comments on the final drafts of the manuscript. The authors wish to thank Prof. Sergio Martinoia, Prof. Sylvie Renaud, Prof. Sylvain Saighi and Prof. Ari Barzilai for their mentorship during the BrainBow project and for useful discussions on the final results.

## Author Contributions

MC, TL, YB, PM, and PB designed the study. YB and TL designed and fabricated the hardware board. SB, VP, FD, and MC designed the experiments. MA, PM, PN, FG and TL designed and worked on SNN. IC, MB, and MT prepared the bimodular cultures. SB and LM performed the experiments. AA and FD performed the lesion experiments. SB, LM, JT, VP, and MC designed the analyses. SB and LM performed the analyses. SB, IC, and VP performed the statistical analyses. SB and IC prepared the original figures. SB, IC, TL, and MC wrote the manuscript. All authors have read, corrected and approved the final version of the manuscript.

## Data and Materials Availability

The datasets generated during and/or analyzed during the current study are available from the corresponding author on reasonable request.

## Supplementary Materials

### Supplementary Information

#### Immunofluorescence Staining and Image Analysis

Immunofluorescence staining was performed as previously described (*75*). Briefly, the cultures on coverslips were fixed with 4% paraformaldehyde in phosphate-buffered saline (PBS) for 30 minutes. After permeabilization with 0.1% Triton X-100 in PBS for 10 minutes four times, the cultures were incubated with PBS containing 5% goat serum and 0.1% Triton X-100 for 1 hour. The permeabilized cultures were incubated with primary antibodies (anti-microtubule associated protein 2 [MAP2] mouse IgG; 1:100; Sigma-Aldrich) in PBS containing 5% goat serum overnight at 4°C and were rinsed with PBS for 10 minutes four times. Then, the cultures were incubated with a secondary antibody (Alexa Fluor 488-labelled anti-mouse IgG; Molecular Probes) in PBS containing 5% goat serum for 2 hours at room temperature and rinsed four times. The coverslips were removed from 12-well plates and mounted on glass slides with mounting media containing DAPI for nuclear staining. Fluorescence images were captured using a fluorescence microscope (Nikon eclips80).

#### SNN Implementation

##### Izhikevich Model

IZH reduces the HH model and proposes an IZH model composed of two equations in which v is the state variable representing the membrane potential of a neuron, and u represents the regeneration of the membrane, which takes into account the activation of the ionic currents K + and the inactivation of the ionic currents Na +.

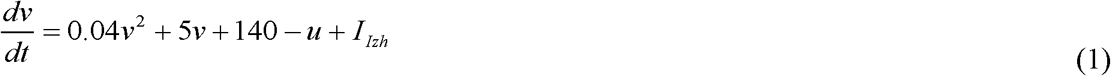

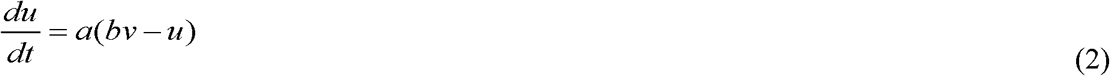

with the after-spike resetting conditions:

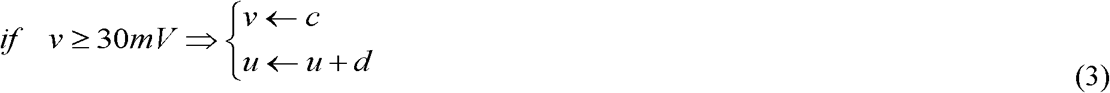

In this model, parameter a describes the time scale of the regeneration variable u(t). Lower values of a indicate slower regeneration. Parameter b describes the sensitivity of the regeneration variable influencing the under-threshold fluctuations of the membrane potential. Parameters c and d are the reset values of the membrane voltage and regeneration variable after an action potential, respectively. The IZH model can reproduce the behavior of all known cortical neurons by changing only 4 parameters (a, b, c and d).

In (3), v is the membrane potential of the neuron, u is a membrane recovery variable that takes into account the activation of potassium and inactivation of sodium channels, and IIzh describes the input current from other neurons.

#### Digital Implementation of the Izhikevich Model

The mathematical equations must be modified to implement the IZH model in digital hardware. To make the IZH neural network more biorealistic, we split the current IIzh into the following currents: Istat, Iexc, and Iinh. Istat is the biasing current, Iexc is the positive contribution due to excitatory synapses and Iinh is the negative contribution of inhibitory synapses.

We also used the methodology developed in (*76*) by multiplying the equation of the membrane voltage by 0.78125 such that all parameters are decomposable with powers of 2, which allows easier digital implementation.

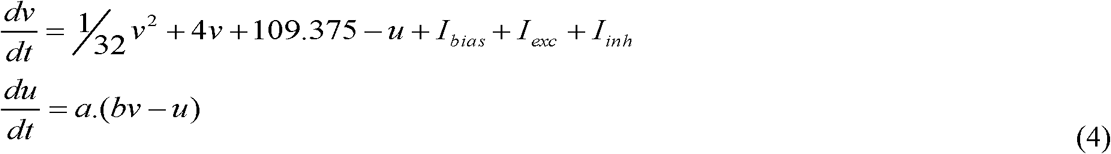

And 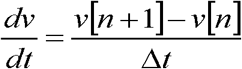 knowing that the computation time of the IZH model is 1 ms

(Δt = 1):

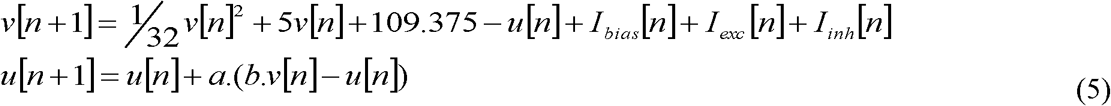

Using a calculation pipeline, we can compute different neurons in parallel. All model parameters are stored in RAM. fig. S3B1 describes the calculation pipelines for v and u.

#### Implementation of Synapse and Synaptic Plasticity

##### Synapse Model

When Wsyn is positive, the synapse is excitatory, and when Wsyn is negative, the synapse is inhibitory. The current Iexc is always positive, and the current Iinh is always negative. However, these two synaptic currents obey the same law of exponential decay. Isyn, the synaptic current, is written:

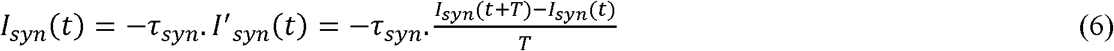

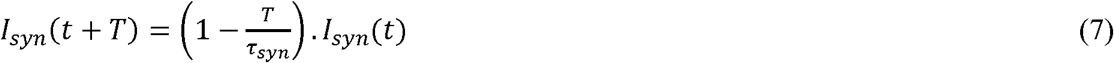

Where computation time T of 1 ms:

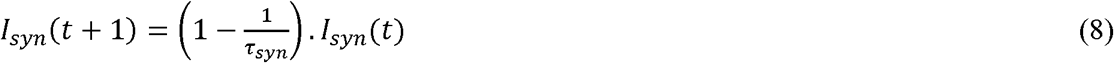

In digital implementation:

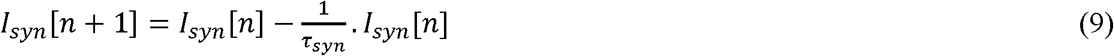

#### Short-Term Synaptic Plasticity Model

This model is defined by 3 parameters:

- a scalar factor xsyn, which indicates the state of the synapse (depression or facilitation) and computes the synaptic weight.
- a percentage P, which will be multiplied by the factor xsyn after each emission of a pre-synaptic action potential. If this percentage is larger than 1, this synapse will describe a short-term facilitation. Otherwise, if this percentage is less than 1, this synapse will describe a short-term depression. In facilitation (or depression), the value added to the stimulation current of a post-synaptic neuron will increase (or decrease) with each emission of action potential.
- a time constant tsyn of exponential decay (or growth) in facilitation (or depression).

When a pre-synaptic spike occurs:

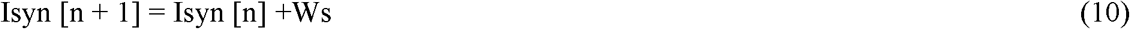

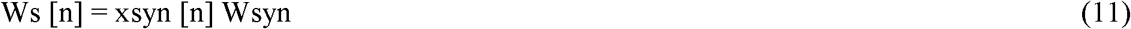

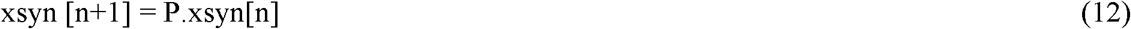

Parameters Wsyn, xsyn, P, τsyn, τampa, and τgaba are stacked into RAM.

#### Synaptic Noise Implementation

The OU process Xt is a prototype of a noisy relaxation process and an example of a Gaussian process that has bounded variance and a stationary probability distribution. The process is stationary, Gaussian and Markovian. This process satisfies the following stochastic differential equation:

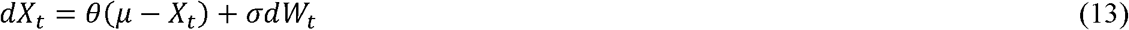

Where *θ*> 0, µ and*σ*> 0 are the parameters, and Wt is the Wiener process.

The parameter μ represents the equilibrium or the mean value of the process. The stationary variance is given by:

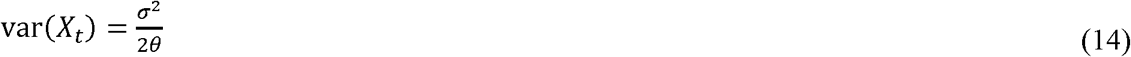

This form of current may represent an approximation to that resulting from the random opening and closing of ion channels on a neuron’s surface or to randomly occurring synaptic input currents with exponential decay (*77*). The digital implementation of this noise in the stimulation current of the neuron is given by:

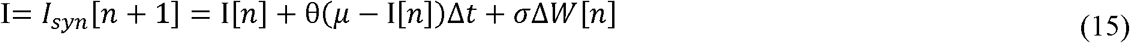

Where n is the iteration step, and Δt = T/N is the time step after the partition of the interval [0, T] into N equal subintervals of width. Note that the random variables ΔW[n] are independent and identically distributed normal random variables with expected value zero and variance Δt, thus ΔW[n]~N(0, Δt) = vΔt * N(0,1).

fig. S5C represents the dynamic behavior of the neuron model in which the parameters are set for tonic bursting activity.

### Online Processing

#### Improving the Signal to Noise Ratio (SNR)

As stated in the main text, the SWT technique we used is composed of a set of orthogonal filters (one highpass, one lowpass). The output of the highpass filter (detail level) was used as an output of the decomposition. The output of the lowpass filter (approximation level) was downsampled, and the filters were applied to provide the next detail and approximation outputs. The process is repeated as long as necessary. From the theoretical point of view, this method split the noise among the different detail outputs but restricted the spike contribution to one or two outputs, which improved the SNR for these specific outputs. The drawback is that the downsampling generated non-stationary behaviors and resulted in sub-optimal detection when the spike time corresponded to dropped samples. To avoid this phenomenon, we chose to replace signal downsampling by filter upscaling (transforming an 8^th^ order filter to a 16^th^ order filter containing 8 interleaved null coefficients). With this technique, the stationary wavelet transform (SWT) (*78*), we kept the best performance for all samples with a price of computational requirements twice as high, and memory requirements 4 times higher. This change in requirements was not an issue since we used hardware computing, and we added the computing resources according to our needs. Although our hardware was able to compute 8^th^ order mother wavelets (highpass/lowpass filter couples), experience showed that the Haar mother wavelet (2^nd^ order) was sufficient [1]. An empirical study showed that the third detail level provides the best performance for detection. As the first detailed output was expected to receive very few contributions from the spikes, it was used to determine the comparison threshold for spike detection.

#### Emphasizing Bursts at the Network Scale

As stated in the main part of our work, the NBD could receive any event coming from the inputs on the system, including ISI-based burst detection events. It was possible to weight the event contributions such that an event coming from an ISI-based detected burst might have a much higher weight than an action potential on a single channel. The 16 NBDs worked on 1 kHz sampled signals because the application they fed was sampled at that frequency. Input events (10 kHz) were combined with a logical OR by group of 10 to produce the downsampling. There was no risk of losing an event because of the behavior of neural cells that were unable to produce two action potentials within 1 ms.

#### Stationary Computing Issue

The NBD detector was very convenient to rapidly check the number of events flowing from the electrodes, but the time slicing resulted in a stationary issue. If number of events able to trigger the BD was produced, the actual detection would depend on whether these events were counted within the same time window or split into two contiguous time windows. As the result of the system relied not only on cell activity but also on the system state at the time the activity occurred, it was non-stationary. This property should be considered when analyzing results. For our application, this property was not considered an issue since the expected network activity bursts are much longer than the computing time window. A stationary detector based on a leaky counter (instead of time windows) was also available on the system to anticipate this behavior. The behavior of such a detector was closer to that of a leaky integrate-and-fire neuron. This variant was not be described because it was not used in the experiments described in this work. Indeed, real-life experiments showed that such detectors were too complex to configure while the cells were being measured.

### Supplementary Figures

**Fig. S1.**
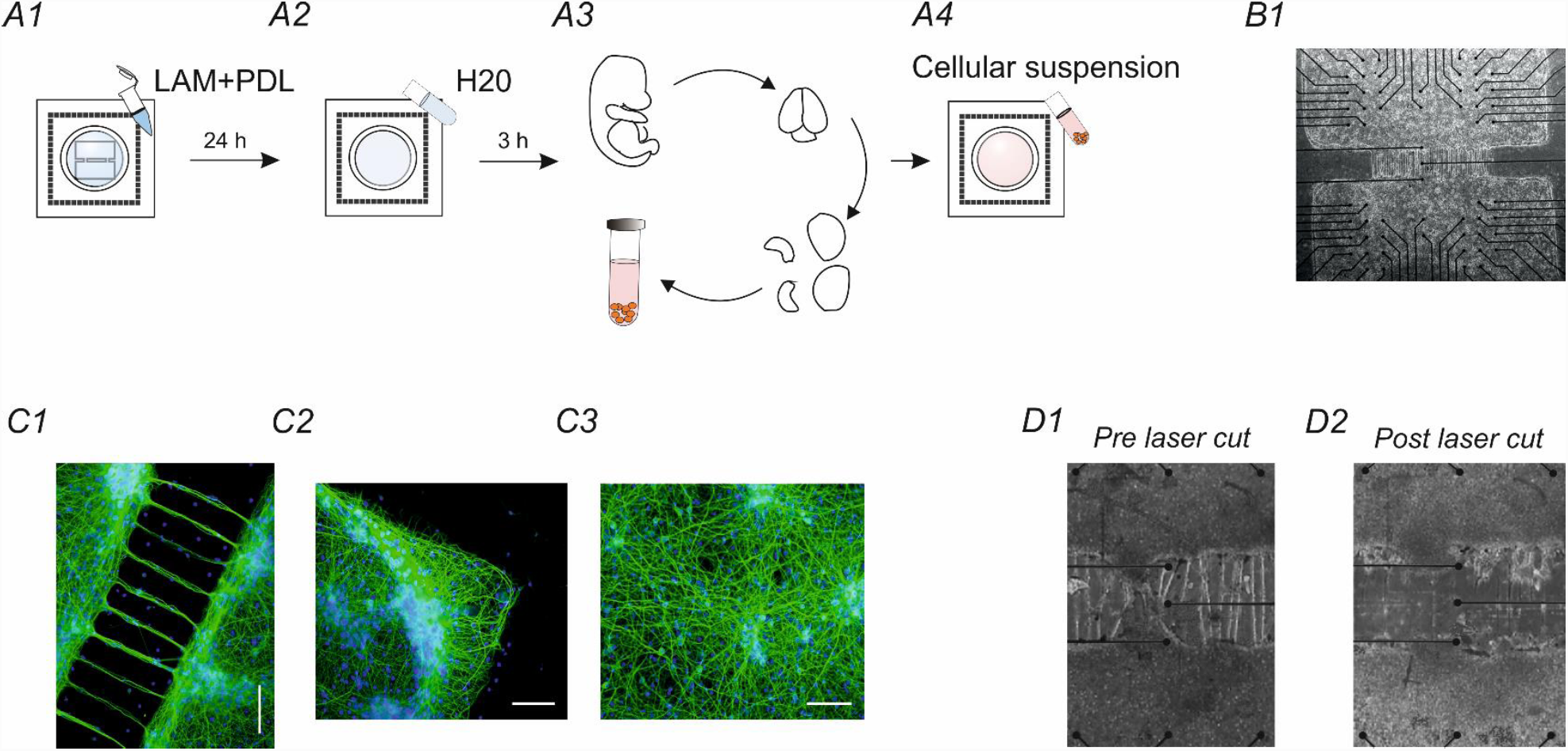
Bimodular neuronal network preparation steps and morphological characterization. (**A1**), MEA surface coating using poly-D-lysine and laminin to promote cell adhesion. (**A2**), Three consecutive surface rinsing steps using sterilized water and 3 hours of waiting before plating cells. (**A3**), Dissection and dissociation procedures of E18 rat neocortex followed by resuspension of the resulting tissue in Neurobasal medium (see ‘Methods’). (**A4**), Cellular solution plating onto a 60-channel MEA surface. (**B1**), A 60-channel MEA (Multichannel Systems MCS, Reutlingen, Germany) with square layout (4Q) plated with a bimodular pattern. (**C1**), Immunofluorescence micrograph of MAP2 of a representative bimodular culture on a coverslip at DIV 25 with magnification of the connections between modules. (**C2**, **C3**), Immunofluorescence micrographs of MAP2 of a representative uniform culture and of one corner on a coverslip at DIV 25. Scale bar = 200 µm. (**D1**), Magnification of the connections between the two modules before performing the laser cut. (**D2**), Magnification of the cut connections between the two modules.

**Fig. S2.**
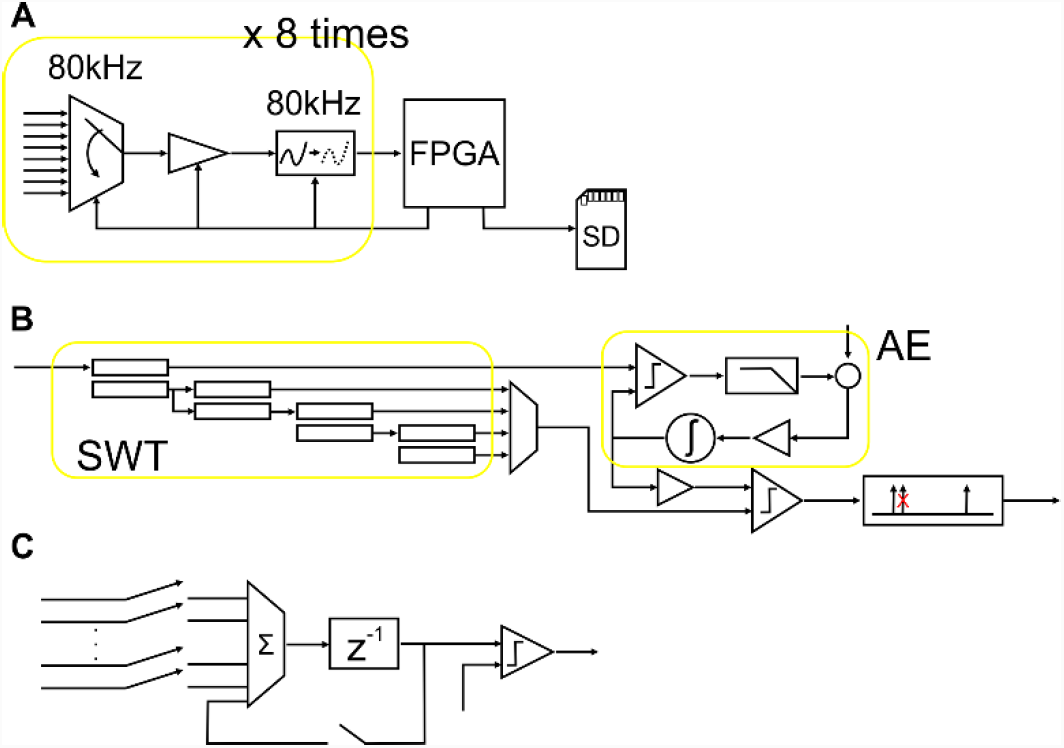
Schematic view of the hardware computation architecture. (**A**), Integrated circuit data flow on the device (Multiplexer: ADG1408, amplifier: LTC6912, analogue to digital converter: AD7686, FPGA: XC6SLX150). Each flow makes it possible to sample 8 channels at 10 kHz (80 kHz multiplexing), eight equivalent structures are connected to the FPGA to sample 64 inputs, and 60 of these inputs are connected to the MEA. (**B**), Hardware spike detection architecture embedded in the FPGA. SWT represents the wavelet-based signal enhancement (stationary wavelet transform). The SWT consists of a series of FIR filters. The first two filters (on the left) are on the 8th order, and the following filters are on the 16th order with 8 null coefficients. AE represents the signal amplitude estimator used to set the final threshold. The loop estimates the standard deviation as the value above 15.9% of samples. The comparator and lowpass filter measure the ratio of samples above the estimated standard deviation; then, the target value is subtracted to identify the relative error. The regulator is composed of an amplifier and integrator to update the estimation of the standard deviation. After the final threshold, an event filter drops each second event too close to the previous one. **c**, Hardware network burst detector, incoming sources are events produced by the spike detector or the ISI-based burst detector, and the event count is stored in the accumulator (z^-1) and cleared according a timer (the accumulator is not actually cleared, but its value is updated without considering the number of accumulated events thus far, which results in data clearance). The number of events counted since the beginning of the time window is compared with the use-defined threshold to decide whether the network is in a burst state.

**Fig. S3.**
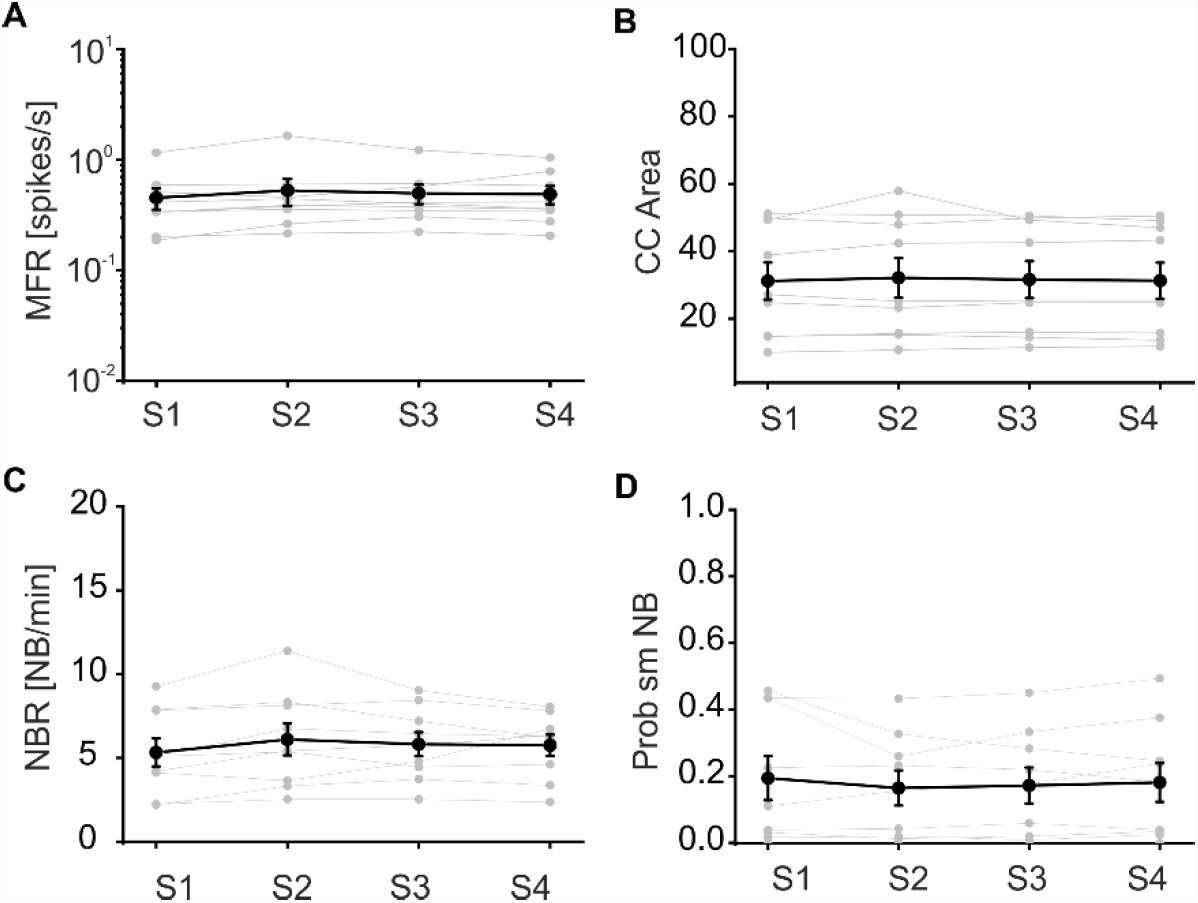
Control no lesion. (**A**), Mean firing rate (MFR) was stable for all control with no lesion experiments for all experimental phases (from S1 to S4). No significant difference was found (Friedman’s repeated measures analysis of variance on ranks; n=9; p=0.04, DF=3, Chi-square=8,333 but no significant difference between rank sums was found by the Tukey test). (**B**), Cross-correlation (CC) area (obtained integrating the CC function of the collapsed spike trains from module 1 and 2 in a range of ± 500 ms) was stable for all control with no lesion experiments for all experimental phases (from S1 to S4). No significant difference was found (Friedman’s repeated measures analysis of variance on ranks; n=9; p=0.833, DF=3, Chi-square=0.867). (**C**), Network burst rate (NBR) was stable for all control with no lesion experiments for all experimental phases (from S1 to S4). No significant difference was found (one-way repeated measures analysis of variance. n=9; p=0.308, DF=3, F=1.267). (**D**), Probability of single-module network burst (Prob smNB) was stable for all control with no lesion experiments for all experimental phases (from S1 to S4). No significant difference was found (Friedman’s repeated measures analysis of variance on ranks; n=9; p=0.789, DF=3, Chi-square=1.050).

**Fig. S4.**
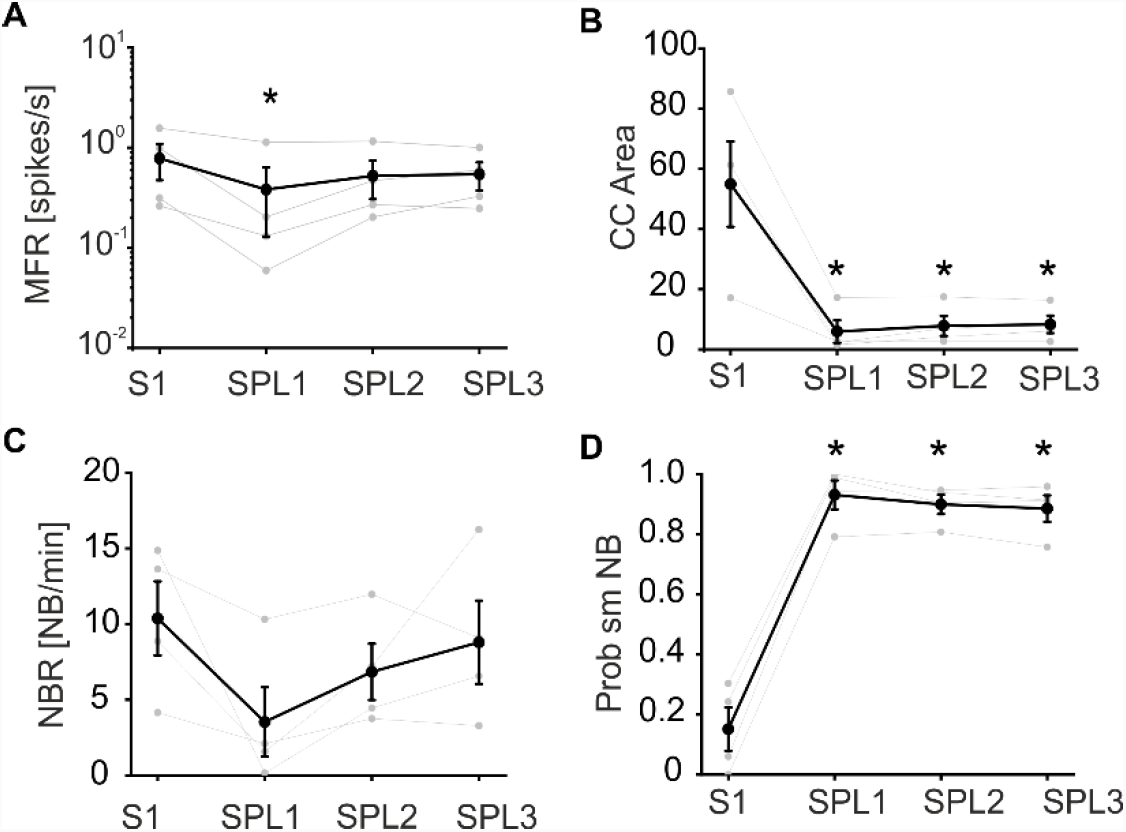
Control with lesion. (**A**), Mean firing rate (MFR) was almost stable for all control with lesion experiments. The only significant difference was found between the first spontaneous phase (S1) before the lesion and the first spontaneous phase post lesion (SPL1). Starting from the second hour post lesion, no significant difference was found between SPL2 and S1 and SPL3 and S1 (one-way repeated measures analysis of variance. n=4; p=0.037, DF=3, F=4,386; all pairwise multiple comparison procedures (Bonferroni t-test): S1 vs. SPL1: p=0.036; S1 vs. SPL2: p=0,282; S1 vs. SPL3: p=0.376; SPL3 vs. SPL1: p=1; SPL3 vs. SPL2: p=1; SPL2 vs. SPL1: p=1). (**B**), Cross-correlation (CC) area (obtained integrating the CC function of the collapsed spike trains from modules 1 and 2 in a range of ± 500 ms) collapsed following laser ablation. We found significant differences between all phases post lesion and the pre-lesion phase (one-way repeated measures analysis of variance. n=4; p<0.001, DF=3, F=16,555; All Pairwise Multiple Comparison Procedures (Bonferroni t-test): S1 vs. SPL1: p=0.001; S1 vs. SPL2: p=0.002; S1 vs. SPL3: p=0.002; SPL3 vs. SPL1: p=1; SPL3 vs. SPL2: p=1; SPL2 vs. SPL1: p=1). (**C**), Network burst rate (NBR) was stable for all control with lesion experiments (from S1 to SPL3). No significant difference was found (one-way repeated measures analysis of variance. n=4; p=0.142, DF=3, F=2.331). (**D**), Probability of single-module network burst (Prob smNB) before the lesion (S1) was on average close to 0.2, meaning that the majority of NBs involved both modules. Following the lesion, the probability increased to an average value higher than 0.85, meaning that the large majority of NBs involved only one module or the other. We found significant differences between SPL1 and S1 (one-way repeated measures analysis of variance. n=4; p<0.001, DF=3, F=109.911; all pairwise multiple comparison procedures (Bonferroni t-test): SPL1 vs. S1: p<0.001; SPL1 vs. SPL3: p=1; SPL1 vs. SPL2: p=1; SPL2 vs. S1: p<0.001; SPL2 vs. SPL3: p=1; SPL3 vs. S1: p<0.001).

**Fig. S5.**
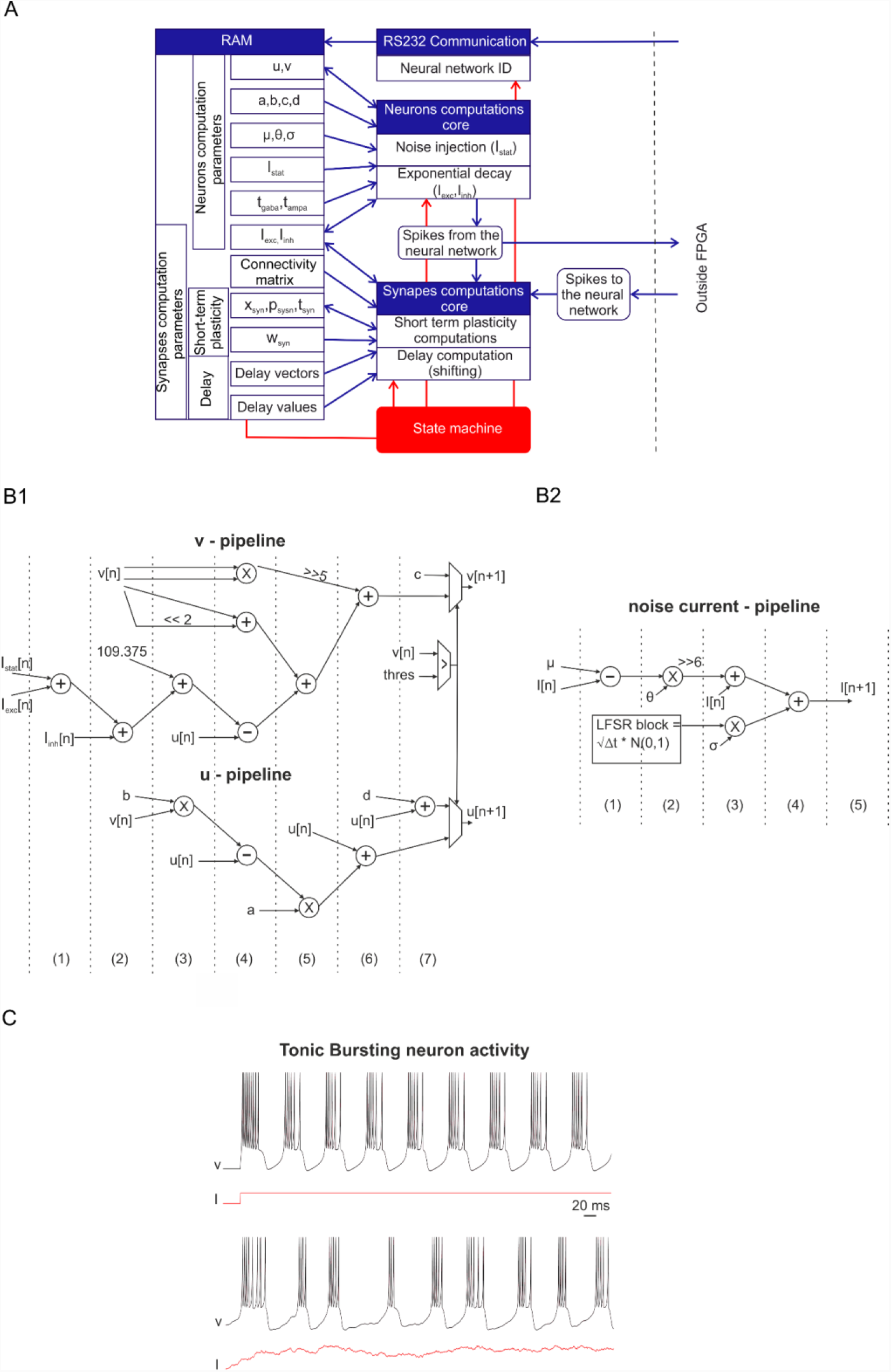
Hardware system organization, pipeline implementation and synaptic noise results. (**A**), Organization of the SNN system with two computation cores, the neuron and synapse. A state machine allows communication between the different blocks. The RAM stacks all the parameters and updates all values every 1 ms. RS232 communication allows communication with a computer to send the initial neural network topology. Spikes to the neural network block are for hybrid experiments where all detected spikes from the neuron culture are sent to the SNN via external synapses. (**B1, B2**), Architecture of ‘v’, and ‘w’ and ‘I’ pipelines for digital implementation. The computation cycles are separated by dotted lines. Each of these dotted lines represents a rising clock edge on which the result of each operation is saved. For the sake of clarity, we have not shown the sequences of the flip-flops. ‘u’ and ‘v’ need 7 clock cycles (140 ns), and ‘I’ needs 5 clock cycles (100 ns). The synaptic noise current ‘I’ is modelled using an Ornstein-Uhlenbeck process, which makes the neuron implementation more biologically plausible, as shown in C. The parameter μ represents the equilibrium or mean value for the process. We set this value equal to 0 with a bias current value necessary for tonic bursting activity without noise. For the other parameters, σ represents the degree of volatility around the mean value caused by shocks, and θ represents the rate by which these shocks dissipate and the variable reverts towards the mean. The stationary variance depends on that parameter. Therefore, in our case, we can set the stationary (long-term) variance with σ and θ parameters (neuron bursting activity) of 35 and 1, respectively. (**C**), SNN output of the tonic bursting neuron in which the parameters are set to obtain tonic bursting activity with and without synaptic noise. The synaptic noise allows for better biomimetic dynamics.

**Fig. S6.**
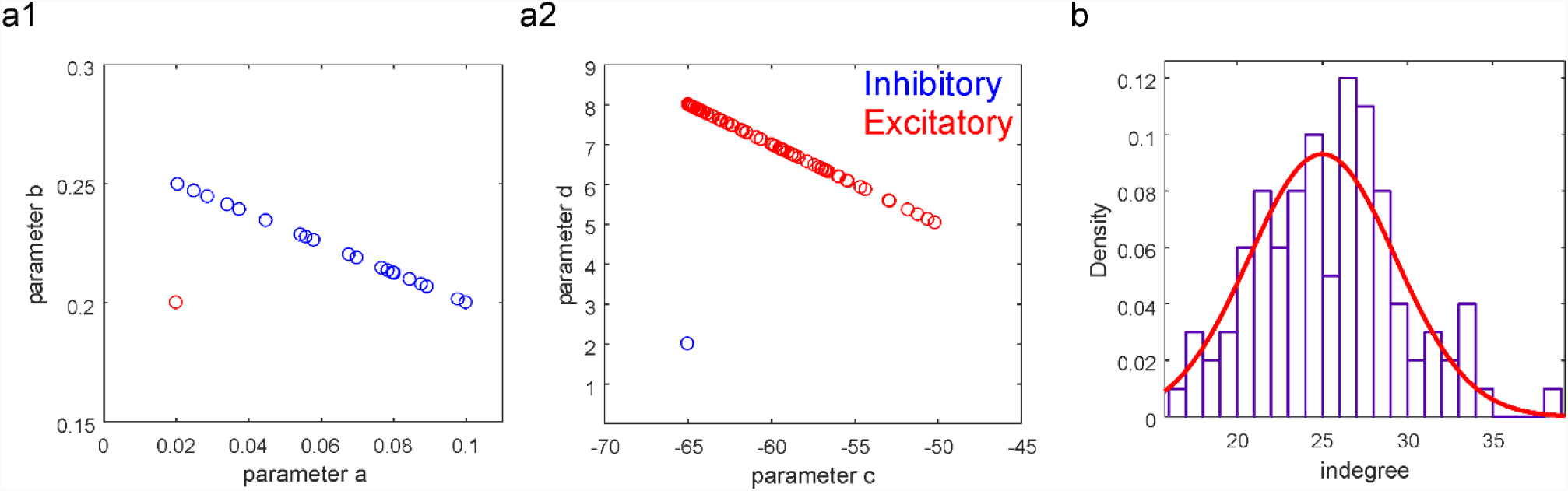
SNN model parameters. (**A1**), Parameters ‘a’ and ‘b’ of the Izhikevich model (see Methods) for excitatory (red circle) and inhibitory (blue circles) neurons. All excitatory neurons have the same parameter a=0.02 and b=0.2. All inhibitory neurons have different parameters a (from 0.02 to 0.1) and b (from 0.2 to 0.25). (**A2**), Parameters ‘c’ and ‘d’ of the Izhikevich model for excitatory (red circles) and inhibitory (blue circles) neurons. All inhibitory neurons have the same parameter c=-65 and d=2. All excitatory neurons have different parameters c (from -64.97 to -50.17) and d (from 5.04 to 7.99). (**B**), Distribution of the indegree (i.e., the number of pre-synaptic neurons) with a Gaussian distribution superimposed (red curve). Mean value = 25, standard deviation = 4.3, R-Square = 0.806 (fit converged, Chi-square tolerance value of 1e-9 was reached).

### Supplementary Tables

**Table S1.** SNN database was created to cover a wide range of NBRs from 1.95 NB/minute (SNN 1) to 94.1 NB/minute (SNN 27). To obtain such variability, we kept the same connectivity matrix with an average degree of 25% for all networks. The main parameters that we tuned to obtain such variability were the mean synaptic excitatory and inhibitory weights. Starting with SNN 1, we added a variable number to each synapse, maintaining separated excitatory (add exc) and inhibitory synapses (add inh). This SNN database can provide a wide range of activities and allows for a better choice of SNN for HBB experiments.

|  | # NB | NB/min | Mean exc | Mean inh | Add exc | Add inh |
| --- | --- | --- | --- | --- | --- | --- |
| **SNN_1** | 39 | 1,95 | 0,99 | -2,02 | 0,00 | 0,00 |
| **SNN_2** | 55 | 2,75 | 1,00 | -2,02 | 0,01 | 0,00 |
| **SNN_3** | 73 | 3,65 | 1,02 | -2,02 | 0,03 | 0,00 |
| **SNN_4** | 90 | 4,50 | 1,03 | -2,02 | 0,04 | 0,00 |
| **SNN_5** | 111 | 5,55 | 1,04 | -2,02 | 0,05 | 0,00 |
| **SNN_6** | 128 | 6,40 | 1,01 | -1,22 | 0,02 | 0,80 |
| **SNN_7** | 136 | 6,80 | 1,00 | -1,02 | 0,01 | 1,00 |
| **SNN_8** | 159 | 7,95 | 1,01 | -1,03 | 0,02 | 0,99 |
| **SNN_9** | 164 | 8,20 | 1,02 | -1,22 | 0,03 | 0,80 |
| **SNN_10** | 186 | 9,30 | 1,02 | -1,04 | 0,03 | 0,98 |
| **SNN_11** | 208 | 10,40 | 1,08 | -2,02 | 0,09 | 0,00 |
| **SNN_12** | 227 | 11,35 | 1,03 | -1,05 | 0,04 | 0,97 |
| **SNN_13** | 239 | 11,95 | 1,08 | -1,92 | 0,09 | 0,10 |
| **SNN_14** | 265 | 13,25 | 1,06 | -1,22 | 0,07 | 0,80 |
| **SNN_15** | 295 | 14,75 | 1,07 | -1,37 | 0,08 | 0,65 |
| **SNN_16** | 303 | 15,15 | 1,08 | -1,52 | 0,09 | 0,50 |
| **SNN_17** | 329 | 16,45 | 1,07 | -1,22 | 0,08 | 0,80 |
| **SNN_18** | 384 | 19,20 | 1,08 | -1,22 | 0,09 | 0,80 |
| **SNN_19** | 420 | 21,00 | 1,09 | -1,22 | 0,10 | 0,80 |
| **SNN_20** | 597 | 29,85 | 1,12 | -1,14 | 0,13 | 0,88 |
| **SNN_21** | 743 | 37,15 | 1,15 | -1,22 | 0,16 | 0,80 |
| **SNN_22** | 842 | 42,10 | 1,16 | -1,18 | 0,17 | 0,84 |
| **SNN_23** | 968 | 48,40 | 1,18 | -1,20 | 0,19 | 0,82 |
| **SNN_24** | 1080 | 54,00 | 1,20 | -1,22 | 0,21 | 0,80 |
| **SNN_25** | 1303 | 65,15 | 1,24 | -1,26 | 0,25 | 0,76 |
| **SNN_26** | 1510 | 75,50 | 1,27 | -1,29 | 0,28 | 0,73 |
| **SNN_27** | 1882 | 94,10 | 1,34 | -1,36 | 0,35 | 0,66 |

**Table S2.** Hardware resource usage for the different functions of the hardware board. The signal processing part includes filters, spike detection, and burst detection (for events from electrodes and from the SNN). The SNN is considered a part by itself, and the environment part includes the necessary engineering to interface the system to the real world. This part includes analogue front-end management, computer interface, VGA display management and SD-card storage management. Bit registers (referenced as flip-flops in digital computing) are used to synchronously store local values or module states. Look-Up tables (LUTs) are asynchronous devices used to implement arbitrary 6-input logic gates. Random Access Memory (RAM) blocks are mostly used to store indexed values (e.g., channel samples for filters, and waiting queues). Multipliers are hard-coded embedded multiplier circuits that are available to increase both performance and density since multiplications are resource-demanding functions in hardware. The high majority of RAM blocks are required by the environment for VGA display and SD-card data buffer. The SNN part uses RAM blocks extensively to address the individual parameters of neurons and synapses. The SNN also requires many hardware multipliers because of the original mathematical formalism. Resources for signal processing are more focused on bit registers and LUTs because of pipeline structures that implement computations structures close to the device that stores data. The environmental cost in terms of resources is heavy, but this cost is constant and should not increase if implementing new signal processing features.

|  | bit registers |  |  |  | LUTs |  |  |  | RAM blocks |  |  |  | multipliers |  |  |
| --- | --- | --- | --- | --- | --- | --- | --- | --- | --- | --- | --- | --- | --- | --- | --- |
|  | numb. | Project | Device |  | numb. | Project | Device |  | numb. | Project | Device |  | numb. | Project | Device |
| Signal processing | 4753 | 31% | 3% |  | 6274 | 31% | 7% |  | 11 | 5% | 4% |  | 3 | 7% | 2% |
| SNN | 3433 | 23% | 2% |  | 5747 | 29% | 6% |  | 82 | 39% | 31% |  | 42 | 93% | 23% |
| Environment | 6993 | 46% | 4% |  | 7996 | 40% | 9% |  | 118 | 56% | 44% |  | 0 | 0% | 0% |
| Total | 15179 | 100% | 8% |  | 20017 | 100% | 22% |  | 211 | 100% | 79% |  | 45 | 100% | 25% |

### Supplementary material

#### Video of the lesion

Video recorded during laser ablation of the connections between the two modules (visible on the left and right side) of a bimodular cell culture. The laser dissection system has been previously described (*18*). Video recorded on November 16, 2016. Duration 18 s. Link: https://drive.google.com/open?id=16uzYJ2T4Ll5quQZ933e7FlUxbCG5Or8Y

## References

1. V. L. Feigin, B. Norrving, G. A. Mensah, Global burden of stroke. Circ. Res. 120, 439–448 (2017).

2. A. I. R. Maas et al., Traumatic brain injury: Integrated approaches to improve prevention, clinical care, and research. Lancet Neurol. 16, 987–1048 (2017).

3. S. Vassanelli, M. Mahmud, Trends and challenges in neuroengineering: Toward "intelligent" Neuroprostheses through brain-"brain inspired systems" communication. Front. Neurosci. 10, 438 (2016).

4. F. D. Broccard, S. Joshi, J. Wang, G. Cauwenberghs, Neuromorphic neural interfaces: From neurophysiological inspiration to biohybrid coupling with nervous systems. J. Neural. Eng. 14, 041002 (2017).

5. S. R. Soekadar, N. Birbaumer, M. W. Slutzky, L. G. Cohen, Brain-machine interfaces in neurorehabilitation of stroke. Neurobiol. Dis. 83, 172–179 (2015).

6. S. N. Abdulkader, A. Atia, M.-S. M. Mostafa, Brain computer interfacing: Applications and challenges. Egypt. Inform. J. 16, 213–230 (2015).

7. S. N. Flesher et al., Intracortical microstimulation of human somatosensory cortex. Sci. Transl. Med. 8, 361ra141 (2016).

8. B. Rosin et al., Closed-loop deep brain stimulation is superior in ameliorating parkinsonism. Neuron 72, 370–384 (2011).

9. C. E. Bouton et al., Restoring cortical control of functional movement in a human with quadriplegia. Nature 533, 247 (2016).

10. F. Kohler et al., Closed-loop interaction with the cerebral cortex: A review of wireless implant technology. Brain Comput. Interfaces 4, 146–154 (2017).

11. J. J. Jun et al., Fully integrated silicon probes for high-density recording of neural activity. Nature 551, 232– 236 (2017).

12. N. A. Steinmetz, C. Koch, K. D. Harris, M. Carandini, Challenges and opportunities for large-scale electrophysiology with Neuropixels probes. Curr. Opin. Neurobiol. 50, 92–100 (2018).

13. D. K. Myers et al., From *in vivo* to *in vitro*: The medical device testing paradigm shift. ALTEX 34, 479–500 (2017).

14. D. O. Hebb, The Organization of Behavior: A Neuropsychological Approach. (John Wiley & Sons, New York, NY, 1949).

15. M. Bisio, A. Bosca, V. Pasquale, L. Berdondini, M. Chiappalone, Emergence of bursting activity in connected neuronal sub-populations. PLoS One 9, e107400 (2014).

16. M. Shein-Idelson, E. Ben-Jacob, Y. Hanein, Engineered neuronal circuits: A new platform for studying the role of modular topology. Frontiers in neuroengineering 4, 10 (2011).

17. P. Bonifazi et al., *In vitro* large-scale experimental and theoretical studies for the realization of bi-directional brain-prostheses. Front. Neural Circuits 7, 40 (2013).

18. F. Difato et al., Combined optical tweezers and laser dissector for controlled ablation of functional connections in neural networks. J. Biomed. Opt. 16, 051306 (2011).

19. J. P. Hayes, E. D. Bigler, M. Verfaellie, Traumatic brain injury as a disorder of brain connectivity. J. Int. Neuropsychol. Soc. 22, 120–137 (2016).

20. A. Pirog et al., Multimed: An integrated, multi-application platform for the real-time recording and sub-millisecond processing of biosignals. Sensors 18, 2099 (2018).

21. D. J. Guggenmos et al., Restoration of function after brain damage using a neural prosthesis. Proceedings of the National Academy of Sciences of the United States of America 110, 21177–21182 (2013).

22. E. M. Izhikevich, Simple model of spiking neurons. IEEE Trans. Neural Netw. 14, 1569–1572 (2003).

23. K. Hayashi, R. Kawai-Hirai, A. Harada, K. Takata, Inhibitory neurons from fetal rat cerebral cortex exert delayed axon formation and active migration *in vitro*. J. Cell Sci. 116, 4419–4428 (2003).

24. P. Bonifazi, M. E. Ruaro, V. Torre, Statistical properties of information processing in neuronal networks. Eur. J. Neurosci. 22, 2953–2964 (2005).

25. F. Grassia, T. Kohno, T. Levi, Digital hardware implementation of a stochastic two-dimensional neuron model. J. Physiol. Paris 110, 409–416 (2016).

26. E. M. Izhikevich, Which model to use for cortical spiking neurons? IEEE Trans. Neural Netw. 15, 1063– 1070 (2004).

27. E. M. Izhikevich, G. M. Edelman, Large-scale model of mammalian thalamocortical systems. Proceedings of the National Academy of Sciences of the United States of America 105, 3593–3598 (2008).

28. D. R. Kipke et al., Advanced neurotechnologies for chronic neural interfaces: New horizons and clinical opportunities. J. Neurosci. 28, 11830–11838 (2008).

29. W. Wang et al., Neural interface technology for rehabilitation: Exploiting and promoting neuroplasticity. Phys. Med. Rehabil. Clin. N. Am. 21, 157–178 (2010).

30. H. A. Johnson, A. Goel, D. V. Buonomano, Neural dynamics of *in vitro* cortical networks reflects experienced temporal patterns. Nat. Neurosci. 13, 917 (2010).

31. G. O. Javier, J. Soriano, E. Alvarez-Lacalle, S. Teller, J. Casademunt, Noise focusing and the emergence of coherent activity in neuronal cultures. Nat. Phys. 9, 582 (2013).

32. S. M. Potter, Closing the loop between neurons and neurotechnology. Front. Neurosci. 4, 15 (2010).

33. S. Kanner et al., Design, surface treatment, cellular plating, and culturing of modular neuronal networks composed of functionally inter-connected circuits. J. Vis. Exp., doi: 10.3791/52572 (2015).

34. H. Yamamoto et al., Impact of modular organization on dynamical richness in cortical networks. Science advances 4, eaau4914 (2018).

35. R. Habibey, A. Golabchi, S. Latifi, F. Difato, A. Blau, A microchannel device tailored to laser axotomy and long-term microelectrode array electrophysiology of functional regeneration. Lab on a chip 15, 4578–4590 (2015).

36. A. Soloperto et al., Modulation of neural network activity through single cell ablation: An *in vitro* model of minimally invasive neurosurgery. Molecules 21, 1018 (2016).

37. D. A. Wagenaar, J. Pine, S. M. Potter, Searching for plasticity in dissociated cortical cultures on multi-electrode arrays. J. Negat. Results Biomed. 5, 16 (2006).

38. A. Luczak, B. L. McNaughton, K. D. Harris, Packet-based communication in the cortex. Nat. Rev. Neurosci. 16, 745–755 (2015).

39. S. Panzeri, C. D. Harvey, E. Piasini, P. E. Latham, T. Fellin, Cracking the neural code for sensory perception by combining statistics, intervention, and behavior. Neuron 93, 491–507 (2017).

40. T. D. Kozai, A. S. Jaquins-Gerstl, A. L. Vazquez, A. C. Michael, X. T. Cui, Brain tissue responses to neural implants impact signal sensitivity and intervention strategies. ACS Chem. Neurosci. 6, 48–67 (2015).

41. T. W. Berger et al., A hippocampal cognitive prosthesis: Multi-input, multi-output nonlinear modeling and VLSI implementation. IEEE Trans. Neural Syst. Rehabil. Eng. 20, 198–211 (2012).

42. E. Greenwald, M. R. Masters, N. V. Thakor, Implantable neurotechnologies: Bidirectional neural interfaces-applications and VLSI circuit implementations. Med. Biol. Eng. Comput. 54, 1–17 (2016).

43. P. R. Roelfsema, A. Holtmaat, Control of synaptic plasticity in deep cortical networks. Nat. Rev. Neurosci. 19, 166–180 (2018).

44. A. V. Cuppone, M. Semprini, J. Konczak, Consolidation of human somatosensory memory during motor learning. Behav. Brain Res. 347, 184–192 (2018).

45. D. J. Ostry, M. Darainy, A. A. Mattar, J. Wong, P. L. Gribble, Somatosensory plasticity and motor learning. J. Neurosci. 30, 5384–5393 (2010).

46. N. Takeuchi, S. Izumi, Rehabilitation with poststroke motor recovery: A review with a focus on neural plasticity. Stroke Res. Treat. 2013, 128641 (2013).

47. H. Markram, The human brain project. Sci. Am. 306, 50–55 (2012).

48. F. Melozzi, M. M. Woodman, V. K. Jirsa, C. Bernard, The virtual mouse brain: a computational neuroinformatics platform to study whole mouse brain dynamics. eNeuro 4, doi:10.1523/ENEURO.0111-1517.2017 (2017).

49. L. E. Osborn et al., Prosthesis with neuromorphic multilayered e-dermis perceives touch and pain. Sci. Robot. 20, eaat3818 (2018).

50. G. Valle et al., Biomimetic intraneural sensory feedback enhances sensation naturalness, tactile sensitivity, and manual dexterity in a bidirectional prosthesis. Neuron 100, 37–45 (2018).

51. D. F. Goodman, R. Brette, The brian simulator. Front. Neurosci. 3, 192–197 (2009).

52. R. M. Pizzi et al., A cultured human neural network operates a robotic actuator. Biosystems 95, 137–144 (2009).

53. A. Bruzzone et al., paper presented at the 37th Annual International Conference of the IEEE Engineering in Medicine and Biology Society (EMBC), Milan, Italy, 25–29 August 2015.

54. Z. Chou et al., Bidirectional neural interface: Closed-loop feedback control for hybrid neural systems. Conf. Proc. IEEE Eng. Med. Biol. Soc. 2015, 3949–3952 (2015).

55. R. Hogri et al., A neuro-inspired model-based closed-loop neuroprosthesis for the substitution of a cerebellar learning function in anesthetized rats. Sci. Rep. 5, 8451 (2015).

56. T. Xu, N. Xiao, X. Zhai, C. P. Kwan, C. Tin, Real-time cerebellar neuroprosthetic system based on a spiking neural network model of motor learning. J. Neural. Eng. 15, 016021 (2018).

57. A. Mata, A. J. Fleischman, S. Roy, Characterization of polydimethylsiloxane (PDMS) properties for biomedical micro/nanosystems. Biomedical microdevices 7, 281–293 (2005).

58. A. M. Taylor, N. L. Jeon, Micro-scale and microfluidic devices for neurobiology. Curr. Opin. Neurobiol. 20, 640–647 (2010).

59. G. M. Whitesides, E. Ostuni, S. Takayama, X. Jiang, D. E. Ingber, Soft lithography in biology and biochemistry. Annual review of biomedical engineering 3, 335–373 (2001).

60. D. B. Weibel, W. R. Diluzio, G. M. Whitesides, Microfabrication meets microbiology. Nature reviews. Microbiology 5, 209–218 (2007).

61. I. Colombi, F. Tinarelli, V. Pasquale, V. Tucci, M. Chiappalone, A simplified *in vitro* experimental model encompasses the essential features of sleep. Front. Neurosci. 10, 315 (2016).

62. F. Difato, L. Schibalsky, F. Benfenati, A. Blau, Integration of optical manipulation and electrophysiological tools to modulate and record activity in neural networks. Int. J. Optomechatronics 5, 191–216 (2011).

63. M. S. Aviv et al., Motility flow and growth-cone navigation analysis during *in vitro* neuronal development by long-term bright-field imaging. J. Biomed. Opt. 18, 111415 (2013).

64. A. Blau, T. Neumann, C. Ziegler, F. Benfenati, Replica-moulded polydimethylsiloxane culture vessel lids attenuate osmotic drift in long-term cell cultures. Journal of biosciences 34, 59–69 (2009).

65. M. Frega et al., Cortical cultures coupled to micro-electrode arrays: A novel approach to perform *in vitro* excitotoxicity testing. Neurotoxicology and teratology 34, 116–127 (2012).

66. D. A. Wagenaar, J. Pine, S. M. Potter, Effective parameters for stimulation of dissociated cultures using multi-electrode arrays. J. Neurosci. Methods 138, 27–37 (2004).

67. A. Quotb, Y. Bornat, M. Raoux, J. Lang, S. Renaud, paper presented at the IEEE International Symposium on Circuits and Systems (ISCAS), Seoul, South Korea, 20-23 May 2012.

68. R. R. Harrison, paper presented at the Proceedings of the 25th Annual International Conference of the IEEE Engineering in Medicine and Biology Society (IEEE Cat. No. 03CH37439), Cancun, Mexico, 17-21 September 2003.

69. M. Ambroise, T. Levi, Y. Bornat, S. Saighi, paper presented at the 2013 47th Annual Conference on Information Sciences and Systems (CISS), Baltimore, MD, USA, 20-22 March 2013.

70. Y. Ben-Ari, R. Khazipov, X. Leinekugel, O. Caillard, J. Gaiarsa, GABAA, NMDA and AMPA receptors: A developmentally regulated ’ménage à trois’. Trends Neurosci. 20, 523–529 (1997).

71. M. Rudolph, A. Destexhe, An extended analytic expression for the membrane potential distribution of conductance-based synaptic noise. Neural Comput. 17, 2301–2315 (2005).

72. M. Chiappalone, A. Vato, L. Berdondini, M. Koudelka-Hep, S. Martinoia, Network dynamics and synchronous activity in cultured cortical neurons. Int. J. Neural Syst. 17, 87–103 (2007).

73. D. Eytan, A. Minerbi, N. Ziv, S. Marom, Dopamine-induced dispersion of correlations between action potentials in networks of cortical neurons. J. Neurophysiol. 92, 1817–1824 (2004).

74. V. Pasquale, P. Massobrio, L. L. Bologna, M. Chiappalone, S. Martinoia, Self-organization and neuronal avalanches in networks of dissociated cortical neurons. Neuroscience 153, 1354–1369 (2008).

75. D. Ito et al., Minimum neuron density for synchronized bursts in a rat cortical culture on multi-electrode arrays. Neuroscience 171, 50–61 (2010).

76. A. Cassidy, A. G. Andreou, paper presented at the IEEE Biomedical Circuits and Systems Conference, Baltimore, MD, USA, 20-22 November 2008.

77. H. C. Tuckwell, F. Y. Wan, J. P. Rospars, A spatial stochastic neuronal model with Ornstein-Uhlenbeck input current. Biol. Cybern. 86, 137–145 (2002).

78. J. Pesquet, H. Krim, H. Carfantan, Time-invariant orthonormal wavelet representations. IEEE Trans. Signal Process. 44, 1964–1970 (1996).

